# Genome-wide CRISPR knockout cell screening platform for the disease vector tick species *Ixodes scapularis*

**DOI:** 10.64898/2026.05.05.721418

**Authors:** Matthew Butnaru, William McKenna, Srishti Goswami, Alejandra Wu-Chuang, Enzo Mameli, Abigail Wilcox, Léopoldine Quennesson, Ah-Ram Kim, Austin Veal, Weihang Chen, Hugo Verzone, Elizabeth A. Lane, Hanna J. Laukaitis-Yousey, Chad Araneo, Nisha Singh, Joao H.F. Pedra, Yanhui Hu, Raghuvir Viswanatha, Norbert Perrimon, Stephanie E. Mohr

**Author notes:** **Corresponding author;** Matthew Butnaru, Norbert Perrimon, Stephanie Mohr.

## Abstract

The black legged tick, *Ixodes scapularis*, is a vector of the bacterium that causes Lyme disease and several other illnesses, including anaplasmosis, babesiosis, and tick-borne encephalitis. Although high-quality genome annotations are available for *I. scapularis*, functional understanding of *I. scapularis* genes is limited. To address this, we developed a platform for genome-wide CRISPR-Cas9 knockout screening in *I. scapularis* cells. To evaluate the platform, we performed a screen to identify genes associated with cellular fitness, and screens for resistance to treatment with copper chloride, Antimycin A, or Destruxin A (DA), a cyclic hexadepsipeptide produced by the pathogenic fungus *Metarhizium anisopliae*. In each case, the screens implicate specific sets of conserved and non-conserved *I. scapularis* genes in relevant cellular functions, providing the first experimental evidence of function for a large set of *I. scapularis* genes. Altogether, in this first-of-its-kind effort for the arthropod subclass Acari, we present an unbiased genome-wide CRISPR-Cas9 knockout cell screening platform, related resources, and datasets that will be broadly useful to efficiently uncover cellular functions of *I. scapularis* genes.

## Introduction

Ticks are obligate blood-feeding ectoparasites and as such, they can transmit bloodborne pathogens to humans, livestock, pets, and wild animals. One species of tick, *Ixodes scapularis*, specifically impacts human health as a vector of *Borrelia burgdorferi,* which causes Lyme disease, and several other microbial pathogens, including *Anaplasma phagocytophilum* (human granulocytic anaplasmosis), *B. miyamotoi* (tick-borne relapsing fever), *Babesia microti* (babesiosis), *Ehrlichia muris eauclairensis* (human ehrlichiosis), and Powassan virus (tick-borne encephalitis) ^1^. Due to the impacts of *I. scapularis* and other ticks, there is an urgent need to better understand tick biology.

Genome sequencing and gene annotation is a critical step towards understanding tick gene function and aids application of other ‘omics approaches. As ‘omics technologies improved over time, genome sequences and gene annotations of increasing quality have become available for the ∼2.1 Gb *I. scapularis* genome ^2–6^. In addition, a partial genome sequence is available for the *I. scapularis* ISE6 cell line ^7^, one of several *I. scapularis* cell lines originally isolated from tick embryos ^8^. Availability of *I. scapularis* gene annotations makes it possible to predict function based on what is known about orthologs in other species. However, related genes can have divergent functions and a large proportion of *I. scapularis* genes do not have orthologs with known functions. Thus, altogether, validated functional information is lacking for a large proportion of *I. scapularis* genes.

The lack of functional information about *I. scapularis* genes can be addressed through the application of functional genomics and other ‘omics technologies *in vivo* and in tick cells. For tick genes with clear orthologs, reverse genetic approaches including RNA interference (RNAi) ^9,10^ and CRISPR-Cas editing ^11–13^ can be used to test ortholog-based predictions of gene function. In addition, we are beginning to learn the transcriptional landscape of some tick cell types, for example through single-cell RNAseq studies of *I. scapularis* blood cells (hemocytes) ^14,15^, information that can further inform gene function. The relatively long lifecycle of *I. scapularis* and difficulty of culturing whole ticks in lab settings are barriers to large-scale application of forward genetic, reverse genetic, and other approaches *in vivo*. Use of continuous cell lines can complement *in vivo* studies. In addition, tick cells can also be modified via transgenesis using the Sleeping Beauty (SB) transposon system ^10^. Despite progress, the field would benefit from the availability of additional methods for study of *I. scapularis* ^11^.

We have established a robust and generalizable pooled CRISPR-Cas9-based approach for functional genomics screening that provides an alternative to lentiviral vectors, which are commonly used for mammalian CRISPR screens ^16^. We first developed the approach for genome-wide knockout (KO) screening in *Drosophila* cultured cells ^17,18^, then extended the approach for use in mosquito cell lines ^19^ and applied it at genome-wide scale in *Anopheles* mosquito cells ^20^. Key to the approach is the engineering of cells that support recombination mediated cassette exchange (RMCE), which is used to introduce single guide RNAs (sgRNAs) into cell genomes, thereby providing a molecular barcode for pooled-format screening ^17–19,21^. Specifically, integration of sgRNAs into the genome via RMCE makes it possible to later amplify integrated sgRNAs by PCR, then compare the relative abundance of sgRNAs using next-generation sequencing (NGS), thereby associating cell phenotypes with genotypes. Once a pooled CRISPR screening platform is established for a given cell line, the platform can facilitate a wide range of cell-based assays. For example, CRISPR screens in human and insect cells have revealed new information on diverse topics, including human host cell-flavivirus interactions ^22,23^, insect hormone signaling ^24^, and insect cell receptors of bacterial toxins ^25,26^.

The availability of gene annotations of increasingly high quality ^5,6^, established cell lines ^8^, and other resources suggested that it might be feasible to develop pooled CRISPR-Cas9 screening in *I. scapularis* cell lines. Here, we report the successful establishment of a genome-wide CRISPR-Cas9 KO cell screening platform for *I. scapularis*, including RMCE+ Cas9+ cell lines, pol III promoters for sgRNA expression, a genome-wide sgRNA library, and added support for *I. scapularis* datasets in our PANGEA gene set enrichment analysis (GSEA) tool ^27^. As proof-of-concept, we show that the *I. scapularis* cell screening platform can be used to identify putative fitness-related genes and to identify genes associated with resistance to treatment with copper chloride (CuCl_2_) or Antimycin A. As expected, orthologs of genes linked to fitness, copper transport, or mitochondrial function were identified in the respective screens. Furthermore, we used the platform to identify genes associated with resistance to treatment with Destruxin A (DA), a cyclic hexadepsipeptide toxin produced by the fungus *Metarhizium anisopliae*, which is pathogenic to insects and ticks ^28,29^. Establishment of a screening platform for *I. scapularis* opens the doors to application of unbiased forward-genetic screening to study a wide range of topics in *I. scapularis* cells, including topics relevant to vector biology and tick control.

## Results

### Molecular genetic toolkit and ‘omics datasets for *I. scapularis* cell studies

Establishing CRISPR-Cas9 KO screening in *Drosophila* and mosquito cells required the availability or development of several molecular genetic methods and resources ^17–20^. To establish a new screening platform, minimal requirements include an available cell line(s), suitable pol II and pol III promoters, methods for transfection and genome engineering, cells engineered for RMCE and expressing Cas9, and design, synthesis, and cloning of a genome-wide sgRNA library. Some of these methods or resources had been previously established, including isolation of *I. scapularis* cell lines ^8^, identification of pol II promoters functional in these cells ^30–33^), and a transposon-based system for generation of transgenic cells ^10^. Moreover, methods for CRISPR-Cas9 editing in *I. scapularis* cells ^34^ and *in vivo* ^12^ have been previously reported. Building on this, we optimized, established, or newly developed the additional methods and resources needed to implement our CRISPR-Cas9 screening system in two *I. scapularis* cell lines, ISE18 and ISE6, which we selected as they are commonly used in the field (ISE6), are more robust than other cell lines (ISE18 and ISE6), and have a higher transfection efficiency than other cell lines (ISE18).

More specifically, we first used the Sleeping Beauty (SB) transposon system ^10^ to engineer ISE6 and ISE18 cells to contain RMCE cassettes with a DsRed or iRFP670 marker (RMCE+ ISE18 DsRed; RMCE+ ISE18 iRFP670; RMCE+ ISE6 DsRed; RMCE+ ISE6 iRFP670) (**Fig. 1A**), then used the SB system again to introduce a Cas9 expression cassette (tagBFP marker) into pools of cells enriched for the RMCE cassette (RMCE+ Cas9+ ISE18 DsRed; RMCE+ Cas9+ ISE18 iRFP670; RMCE+ Cas9+ ISE6 DsRed; RMCE+ Cas9+ ISE6 iRFP670) (**Fig. 1B**). In all cases, fluorescence activated cell sorting (FACS) was used to enrich marker-positive (RMCE+ cells) or double marker-positive cell pools (RMCE+, Cas9+ cells). As part of this work, we compared the strengths of several exogenous pol II promoters in ISE6, ISE18, and additional *I. scapularis* cell lines (**Suppl. Fig. 1 A-E**). Furthermore, after using SB to introduce RMCE cassettes into ISE6 and ISE18 cells, we confirmed that the PhiC31 integrase system in general, and RMCE specifically, can function in these cells (RMCE+ ISE18 DsRed, **Fig. 1C,D**; RMCE+ ISE6 DsRed, **Suppl. Fig. 1G**). As expected, when a plasmid expressing PhiC31 is co-transfected with a compatible attB-GFP-attB cassette plasmid (pLib-GFP), we observe a population of cells that remains positive for the GFP marker after transient GFP expression has been lost in control cells (RMCE+ ISE18 DsRed, **Fig. 1D**; RMCE+ ISE6 DsRed**, Suppl. Fig. 1G**).

**Figure 1:**
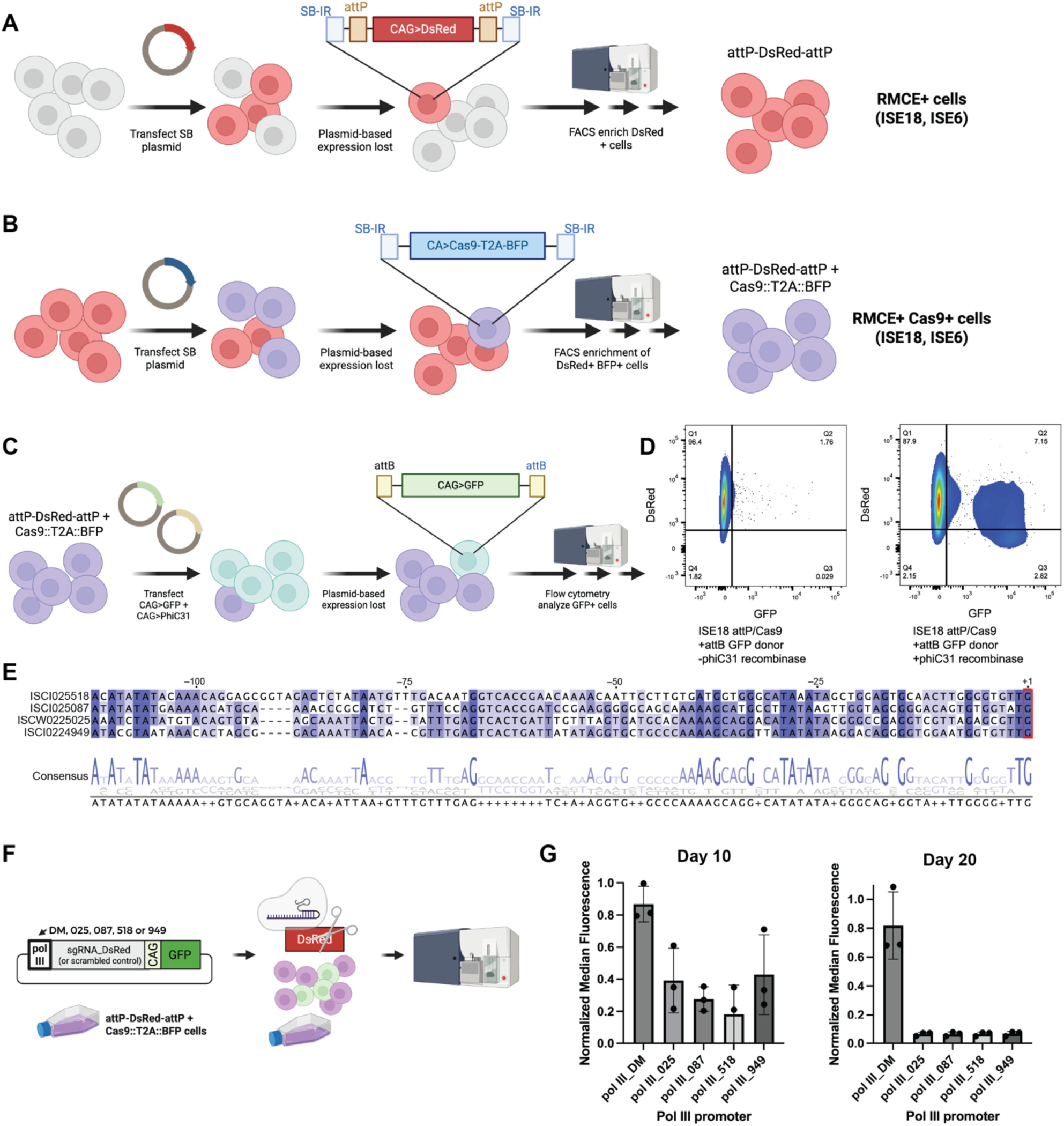
Engineering of *I. scapularis* ISE6 and ISE18 cells to support CRISPR-Cas9 KO screening. (A) – (B) Schematic of the engineering workflow. (A) To establish *I. scapularis* ISE6 and ISE18 cell populations competent for pooled-format CRISPR-Cas9 screening, i.e., RMCE+ Cas9+ cell lines, we used a Sleeping Beauty (SB) to integrate an RMCE cassette in which DsRed is flanked by attP sequences in the genome. (B) The same approach was used to integrate a Cas9-T2A-BFP plasmid into the RMCE+ cells. (C) Design of assay for RMCE activity in RMCE+ cells. (D) Example of FACS data following RMCE in RMCE+ ISE18 DsRed cells in the absence (left) or presence (right) of a plasmid source of PhiC31 integrase. In the presence of integrase, GFP expression is retained over time in some cells, consistent with RMCE-based integration of the GFP cassette. (E) Alignment and consensus for putative *I. scapularis* pol III promoter regions. Sequences included in Materials (**Suppl. File 12**). VectorBase IDs are indicated. Red, putative transcription start site. (F) Design of assay used to test candidate pol III promoters and confirm Cas9 activity. If the pol III promoter results in expression of the sgRNA and Cas9 is active, then reduced levels of DsRed should be observed in GFP-positive cells. DM, *D. melanogaster* pol III promoter; labeling for *I. scapularis* candidates based on VectorBase IDs. (G) Pol III activity assay in RMCE+ Cas9+ ISE18 DsRed cells. Quantification of DsRed fluorescence at day 10 (left) and day 20 (right).

We next sought to confirm Cas9 activity in these cells by transfecting sgRNAs in a manner compatible with pooled screening workflows. For screens, we deliver sgRNA expression cassettes to RMCE-competent cells in plasmid form, with attB sites flanking a pol III promoter-driven sgRNA module that allows for inexpensive and straightforward cloning of sgRNAs into the plasmid. To build sgRNA expression cassette plasmids for *I. scapularis*, we first identified putative pol III promoters bioinformatically based on matches to the U6 promoter sequence upstream of U6-type small nuclear RNA (snRNA) genes annotated in the *I. scapularis* genome, similar to the approach we used to identify pol III promoters for use in mosquito cell lines ^19^ (**Fig. 1E**). Then, we used CRISPR-Cas9-based editing of the DsRed marker present in our engineered RMCE+ Cas9+ cells as an indirect indicator of sgRNA expression (RMCE+ Cas9+ ISE18 DsRed, **Fig. 1F,G**); RMCE+ Cas9+ ISE6 DsRed, **Suppl. Fig. 1F**. Notably, for this analysis, we did not co-transfect a source of integrase, such that disruption of DsRed could only result from expression of the DsRed-targeting sgRNA and subsequent Cas9 editing. Using this assay, we found that by day 10, we observe reduced levels of DsRed when the DsRed-targeting sgRNA is under the control of any of the four candidate *I. scapularis* promoters, suggestive of sgRNA expression and KO, but not with the *D. melanogaster* pol III promoter (**Fig. 1G, Suppl. Fig. 1F**). We note that one of these pol III promoter plasmids was used by our group in a study that included CRISPR-Cas9 over-expression ^15^. To design a genome-wide library for KO and facilitate other CRISPR-Cas9 studies in *I. scapularis*, we precomputed Cas9-compatible sgRNA designs for the *I. scapularis* reference genome (NCBI reference genome annotation ASM1692078v2) and made the resulting genome-wide *I. scapularis* sgRNA designs searchable and downloadable at our previously reported CRISPR GuideXpress online resource ^19^.

We next performed RNAseq in ISE18, ISE6, and a third cell line, IDE2, and compared their profiles to obtain data useful for later screen data quality analysis. The ISE18 and ISE6 expression profiles are highly correlated, whereas the IDE2 cell line is more divergent (**Suppl. Fig. 2**). As part of this analysis, we compared ISE18 wildtype cells with ISE18 cells modified with the DsRed RMCE expression cassette (RMCE+ ISE18 DsRed) and found that the wildtype and engineered cell lines have nearly identical transcriptomes (**Suppl. Fig. 2**). Altogether, we applied, optimized, or newly established all the components needed for application of our CRISPR-Cas9 KO screening approach in *I. scapularis* ISE18 or ISE6 cells, including production of enriched pools of engineered RMCE+ Cas9+ cells, sgRNA expression plasmids with functional pol III promoters, a genome-wide set of sgRNA designs, and transcript and genome datasets useful for data quality analysis. We moved forward in next steps with the engineered ISE18 cells because we found them to be more robust and easier to transfect than ISE6 cells.

### Genome-wide screen identifies high-confidence fitness genes in *I. scapularis* cells

To test the newly established platform, we first selected a subset of 150,972 sgRNA designs targeting 24,340 *I. scapularis* genes, then filtered the list to exclude 9,054 genes not expressed in ISE18, ISE6, or IDE2 cell lines based on our RNAseq data. We synthesized the resulting ∼100K sgRNA library along with 100 scrambled controls and cloned them into a pol III expression cassette compatible with the RMCE system for genome integration, resulting in a “version 1.0” *I. scapularis* sgRNA library (for a summary of gene coverage, see **Table 1**). Next, we transfected the plasmid library into an enriched pool of RMCE+ Cas9+ ISE18 DsRed cells along with the PhiC31 expression plasmid, followed by two rounds of FACS enrichment for the fluorescent marker in the sgRNA cassette over eight weeks of outgrowth (workflow outlined in **Suppl. Fig. 3**). We included two biological replicates (distinct library plasmid preps and transfections), one with two technical replicates and another with one, and report data for all three replicates in **Suppl. File 1**. For each replicate, we PCR amplified integrated sgRNAs and performed NGS. Subsequent to library synthesis, we sequenced the ISE18 cell genome, identified SNPs in the cell genome as compared with the reference sequence, and filtered out sgRNAs that have SNPs, resulting in a virtual library (v1.1; **Table 1**). Only sgRNAs remaining in the v1.1 library were used for screen data analysis (see **Methods** and **Suppl. File 1).** Genes that drop out are likely to be genes required for cell viability and/or growth, or otherwise required for cell fitness (hereafter, “fitness genes”; **Fig. 2A**). Genes enriched in the KO cell pool are likely to encode proteins that restrict or limit growth (hereafter, “growth-restrictive genes”; **Fig. 2B**).

**Figure 2:**
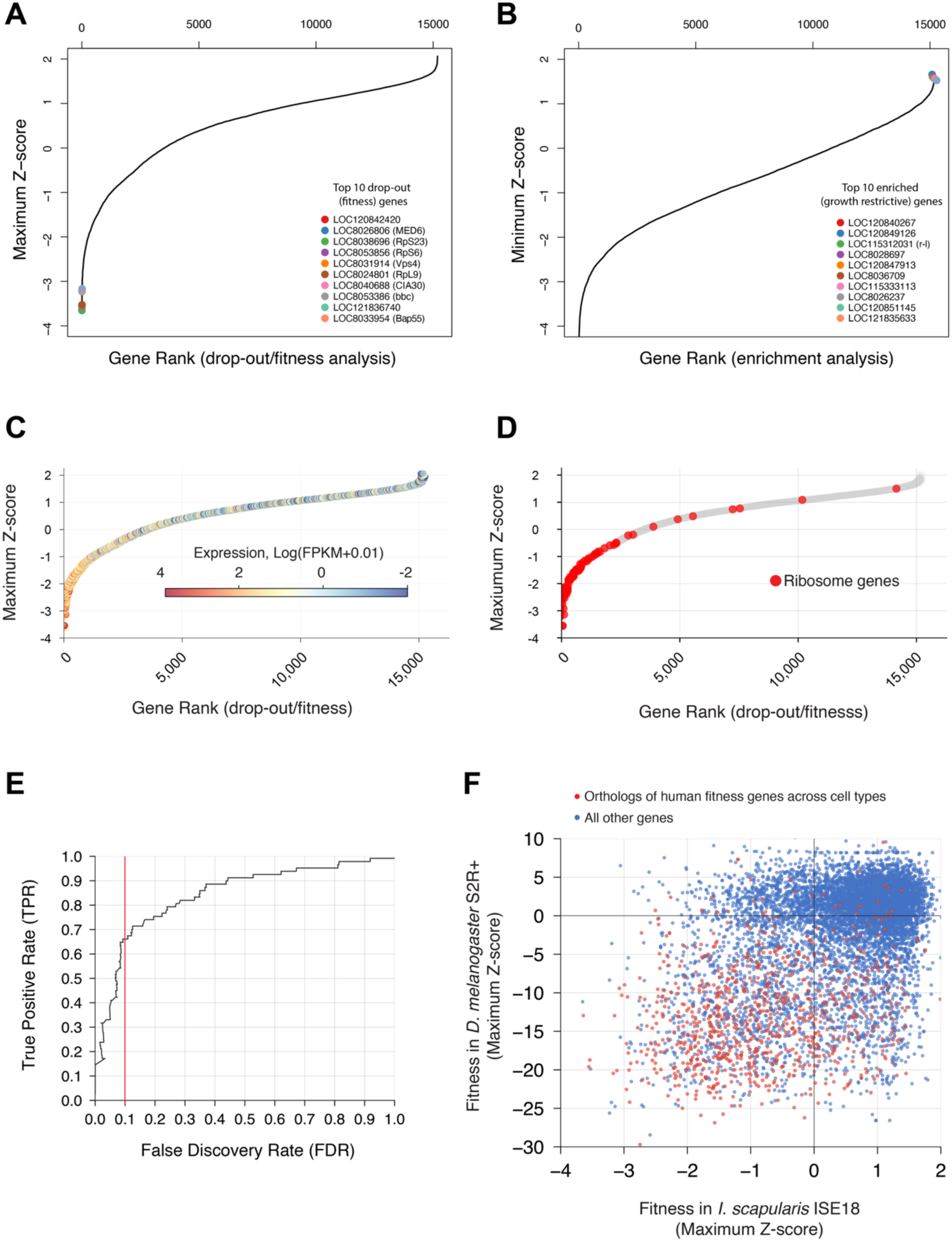
Genome-wide ‘drop-out’ screen identifies *putative I. scapularis* fitness genes and growth-restrictive genes. (A) Plot of maximum Z-score from three replicate drop-out assays in ISE18 cells. The ten top-scoring putative fitness genes are indicated by systematic identifier (gene symbols in parenthesis). (B) Plot of minimum Z-score from three replicates. The ten top-scoring putative growth-restrictive genes are indicated (gene symbol in parenthesis). (C) Maximum Z-scores (fitness scores) with expression levels indicated by heatmap. D) Maximum Z-scores (fitness scores) with genes annotated to encode components of the ribosome indicated in red. (E) Plot of false discovery rate (FDR) versus true positive rate (TPR). FDR estimated based on gene expression; TPR estimated based on ribosomal genes. Red line, cutoff of FDR 0.1 used for GSEA. (F) Plot of maximum Z-score (fitness score) for *I. scapularis* genes with orthologs in *D. melanogaster* versus maximum Z-score (fitness score) for the *D. melanogaster* orthologs of those genes, as determined using a similar screen in S2R+ cells. Red, genes that have human orthologs that score as fitness genes across multiple screens as reported in DepMap. Blue, all other genes.

**Table 1:** Coverage of *I. scapularis* genes in the sgRNA library.

| # of sgRNAs per gene | full coverage designs |  | v1.0 library (screening library<sup>1</sup>) |  | v1.1 SNP-filtered library (data analysis library<sup>2</sup>) |  |
| --- | --- | --- | --- | --- | --- | --- |
|  | # of genes | # of sgRNAs | # of genes | # of sgRNAs | # of genes | # of sgRNAs |
| 1 | 1,060 | 1,060 | 361 | 361 | 458 | 458 |
| 2 | 1,074 | 2,148 | 389 | 778 | 560 | 1,120 |
| 3 | 905 | 2,715 | 347 | 1,041 | 840 | 2,520 |
| 4 | 710 | 2,840 | 270 | 1,080 | 1,534 | 6,136 |
| 5 | 710 | 3,550 | 291 | 1,455 | 2,928 | 14,640 |
| 6 | 608 | 3,648 | 258 | 1,548 | 4,203 | 25,218 |
| 7 | 19,273 | 134,911 | 13,370 | 93,590 | 4,663 | 32,641 |
| Control sgRNAs |  | 100 |  | 100 |  | 100 |
| Total | 24,340 | 150,972 | 15,286 | 99,953 | 15,186 | 82,833 |

<sup>1</sup> Coverage of genes not expressed in any cell line tested in our RNAseq experiment were removed from the full-coverage design set prior to synthesis, resulting in the v1.0 library.
<sup>2</sup> Genome sequence data for ISE18 cells was used to remove sgRNAs with SNPs in the cell line as compared with the reference genome, resulting in the virtual v1.1 library, which was used for screen data analysis.

We next sought to more rigorously assess the accuracy of the screen in determining which genes cause fitness defects upon KO. To do this, we estimated the false discovery rate (FDR) using our ISE18 gene expression data, applying the logic that genes essential for optimal cell fitness must be actively transcribed. We first asked if, as expected, expressed genes were enriched among fitness genes, and found a strong correlation (**Fig 2C**). Since ribosome genes are highly conserved and universally cell-essential in most organisms, we next asked whether ribosome genes would serve as a reliable set of true positives and found them to be highly enriched among the top 2,000 fitness genes (**Fig. 2D**). Using these benchmarks, we then quantified screen accuracy by measuring the rate at which non-expressed genes were falsely recovered as fitness genes (false discovery rate, FDR) relative to the rate at which ribosome genes (true positives) were identified (**Fig. 2E**). We find that the screen has a recall of approximately 65% of true positives at FDR 0.1. As an orthogonal measure of screen accuracy, we asked whether other known functional gene categories were similarly enriched among fitness genes using gene set enrichment analysis (GSEA). We used the FDR cutoff value of 0.1 to define a set of 850 *I. scapularis* fitness genes for GSEA. To directly perform GSEA with *I. scapularis* gene information, we modified our PANGEA online resource ^27^ to include the gene ontology annotations for *I. scapularis*. We then used PANGEA to analyze the set of 850 fitness genes. As expected, we see statistically significant enrichment for fundamental biological processes such as translation, mRNA splicing, and transcription initiation and fundamental molecular functions such as ATP binding, RNA binding, and GTPase activity (**Suppl. File 2**).

Additionally, we wanted to take advantage of extensive functional annotation of *D. melanogaster* to further analyze the set of *I. scapularis* fitness genes. To facilitate this, we first mapped orthologs at genome scale using our DRSC Integrative Ortholog Prediction Tool (DIOPT) approach ^35^ among *I. scapularis, D. melanogaster,* human, and other organisms, and expanded DIOPT online resources to include the *I. scapularis* ortholog mapping to other organisms ^36^. Application of the DIOPT approach identified orthologs for 8,791 of the 15,287 *I. scapularis* genes screened (58%). In the subset of *I. scapularis* fitness genes (FDR cutoff 0.1), DIOPT identified *D. melanogaster* orthologs for 773 of 850 (91%) (**Suppl. File 1**). We again used PANGEA to perform GSEA, this time using the set of *D. melanogaster* orthologs of the *I. scapularis* fitness genes as the query list, and the full set of 8,791 orthologs of *I. scapularis* genes screened for background correction. As expected, we found strong enrichment for gene sets related to fundamental biological processes (e.g., “cytoplasmic translation”; 4.9-fold enrichment, P-value = 1E-25) and molecular functions (e.g., “structural constituent of ribosome”; 3.8-fold enrichment, P-value = 1.26446E-24), and for FlyBase annotated phenotypes including “cell lethal” (3.1-fold enrichment, P-value = 7.906E-07) (**Suppl. File 3**).

We were also interested in broader conservation of fitness genes. To address this, we performed cross-species comparison of fitness factors as identified in similar screens in *D. melanogaster* or human cell lines. As expected, a large fraction of genes identified as fitness genes in *I. scapularis* have orthologs that were identified as fitness genes in a CRISPR KO cell screen with *D. melanogaster* S2R+ cells ^37^ (**Fig. 2F**). Specifically, of the 773 *I. scapularis* fitness genes (FDR < 0.1) with *D. melanogaster* orthologs, 632 (82%) were also fitness genes in S2R+ cells (FDR < 0.05), more than twice the background rate of 34% across all orthologous genes (hypergeometric test, p < 1E-174, Bonferroni-corrected; **Fig. 2F**, **Suppl. File 1**). Similarly, of the 794 *I. scapularis* fitness genes with human orthologs, 316 (40%) corresponded to human fitness genes as defined by whole-genome CRISPR screens in human cells ^38^, four times the background rate of 10% across all orthologous genes (hypergeometric test, p < 1E-146, Bonferroni-corrected; **Fig. 2F, Suppl. File 1**).

To begin to explore differences between fitness requirements *I. scapularis* and *D. melanogaster*, we also performed GSEA with the subset of *I. scapularis* fitness genes that have orthologs in *D. melanogaster* that did not score as fitness genes in S2R+ cells. We see statistically significant enrichment in this subset of genes for transforming growth factor-beta/bone morphogenic protein (TGFb/BMP) signaling pathway components, including enrichment for the biological process terms “transforming growth factor beta receptor superfamily signaling pathway” “BMP signaling pathway,” “response to BMP,” and “positive regulation of SMAD protein signal transduction,” suggesting that *I. scapularis* ISE18 cells are dependent on TGFb/BMP signaling (**Suppl. File 4**). Three canonical TGFb/BMP signaling pathway components score as fitness genes in the screen, LOC115309488 (*I. scapularis* gene description “bone morphogenetic protein receptor type-1B”; ortholog of *D. melanogaster thickveins*), LOC8036103 (bone morphogenetic protein receptor type-2; ortholog of *wishful thinking*), and LOC8038463 (TGF-beta receptor type-1; ortholog of *baboon*). Because the screen can detect only cell-autonomous fitness factors, we do not expect to see enrichment for a ligand. However, consistent with the idea that the pathway is active in ISE18 cells, we do observe that one of the *I. scapularis* genes that encodes a TGFb/BMP-type ligand is expressed at appreciable levels in ISE18 cells, LOC8031831 (bone morphogenetic protein 7, ortholog of *glass bottom boat;* FPKM 20 in *I. scapularis* ISE18 cells; see **Suppl. File 5**).

Regarding putative growth-restrictive genes, as might be expected for a continuous cell line, there are no strong statistical hits. Nevertheless, PANGEA analysis of a set of hits defined by applying a loose cutoff (minimum enrichment score ≥ to 1.3) reveals significant enrichment for the biological processes “tRNA wobble uridine modification” (50-fold enrichment, P-value = 0.000694768) and “glutathione metabolic process” (11-fold enrichment; P-value = 0.002821417), as well as for the molecular function “glutathione transferase activity” (11-fold enrichment; P-value 0.002339738) and other functions (**Suppl. File 6**). GSEA with the *Drosophila* orthologs of the same list reveals enrichment for “Positive Regulators of Toll-NF-kappaB Signaling Pathway” (12-fold enrichment, P-value = 0.011499982). These results suggest that Toll signaling might be growth restrictive in ISE18 cells. Consistent with this, an ortholog of *Pellino (Pli)*, which acts as a negative regulator of Toll signaling in *D. melanogaster* ^39^, is among the top 100 genes identified as *I. scapularis* fitness genes (LOC8035299; rank = 63; enrichment scores −2.8, −2.6, −3.5 in the three replicates; see **Suppl. File 1**). The idea that Toll signaling can be growth restrictive is consistent with findings for *D. melanogaster* ^40^.

### Knockout of *I. scapularis Ctr1A* confers resistance to treatment with copper chloride

We next sought to establish a proof-of-concept selection assay, as selection assays were useful for developing and validating screen platforms in *Drosophila* and mosquito cells ^18,19^ and selection-based screens have been particularly powerful for uncovering novel gene functions in *Drosophila* and mosquito cells (e.g., see ^24–26,41,42^). To establish a selection assay, we sought a treatment that was toxic to ISE18 cells and for which we could predict that KO of a specific *I. scapularis* gene would result in resistance to that treatment. Supplementation of cell culture media with metal salts is toxic to insect cells and knockdown of ion transporters can confer resistance to the treatment ^43^. We found that CuCl_2_ supplementation of *I. scapularis* culture media is toxic to ISE18 cells. Moreover, the *I. scapularis* gene annotation includes *Ctr1A* (LOC120838508), a putative ortholog of *Drosophila Ctr1A*, which has a predicted “Ctr copper transporter” domain (InterPro ID IPR007274) ^44^ and is known to function as a copper transporter ^45^.

To further compare *I. scapularis* Ctr1A with well-characterized copper transporters, we next analyzed the predicted Ctr1A structure. Specifically, to determine the 3D structure and optimal stoichiometry of Ctr1A, we used AlphaFold 3 ^46^ to predict oligomeric states of *I. scapularis* Ctr1A ranging from 2 to 5 subunits and evaluated each using the integrated Local Interaction Score (iLIS) and ipTM values ^47^ (**Suppl. Fig. 4A**). The trimeric configuration yielded the highest iLIS and ipTM scores among all tested stoichiometries (**Fig. 3A; Suppl. Fig. 4A**), suggesting that the homotrimer is the most stable oligomeric state, consistent with what is known for the human CTR1 family ortholog ^48^. To assess ion binding selectivity, we predicted the Ctr1A trimeric complex in the presence of each ion species available in the AlphaFold 3. Cu²⁺ produced the highest iLIS and ipTM scores, followed by Co²⁺ and Fe²⁺, while all other ions showed substantially lower scores (**Fig. 3B,C; Suppl. Fig. 4B**). Furthermore, comparison of Ctr1A trimer predictions with and without Cu²⁺ revealed that inclusion of Cu²⁺ increased the overall predicted structural confidence (pLDDT) (**Suppl. Fig. 4C,D**), suggesting that copper binding stabilizes the complex. In the predicted structure, the Cu²⁺ ion was positioned at the entrance of the central channel of the trimeric pore, coordinated by methionine residues (M203 and M207; **Fig. 3A-C**). Altogether, these results suggest that *I. scapularis Ctr1A* encodes a canonical CTR1 family copper transporter that can form a homotrimeric complex with preferential binding toward Cu²⁺.

**Figure 3:**
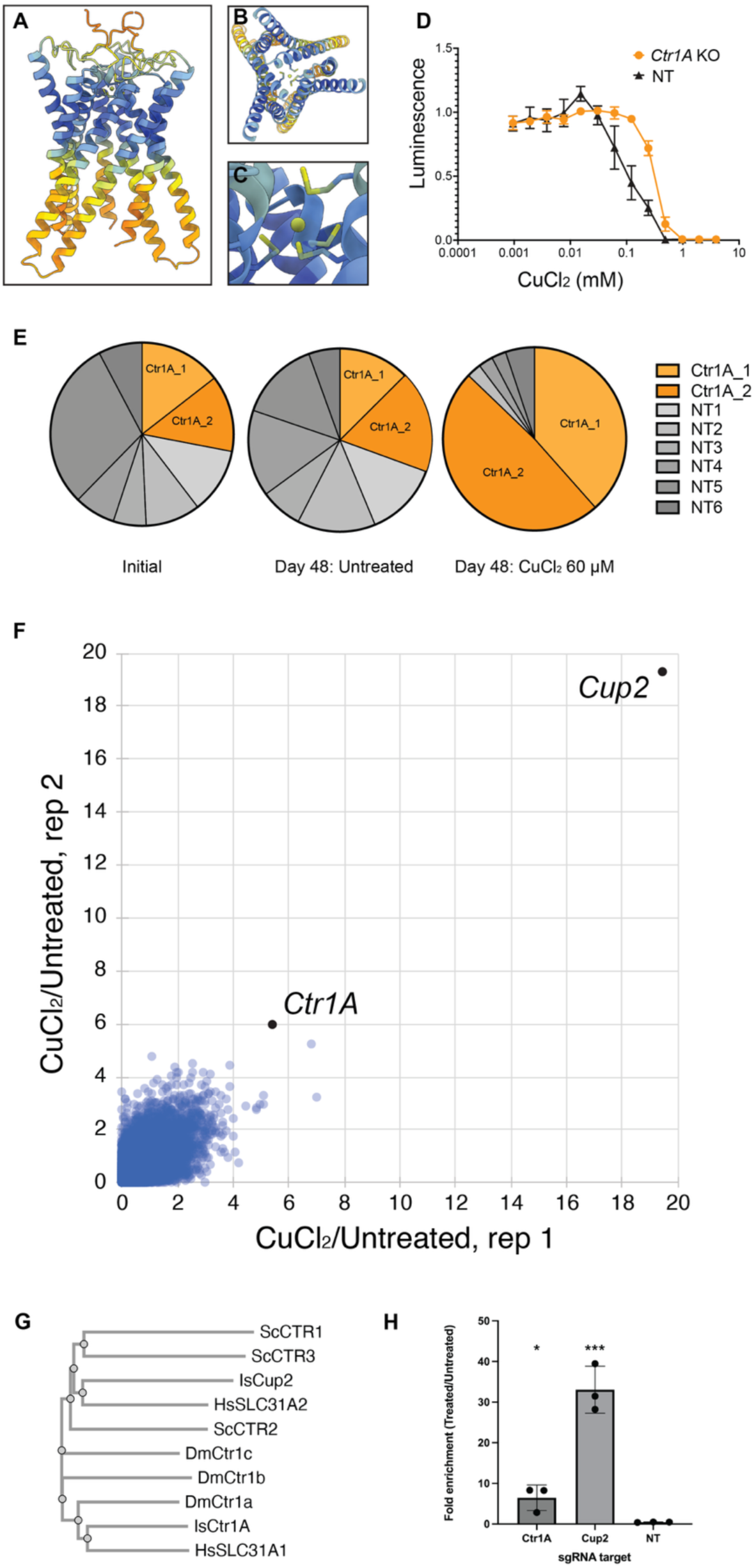
Genome-wide screen for CuCl_2_ resistance identifies gene encoding CTR1 family copper transporters. (A) – (C), AlphaFold3 predicted 3D model of an *I. scapularis* Ctr1A trimeric complex with a copper ion (sphere) bound in a central pore. (A), side view; (B) top view, (C), close-up view of the copper ion binding region. (D) Total ATP assay comparing cells transfected with a Ctr1A-targeting sgRNA versus with a non-targeting sgRNA and treated with CuCl_2_. Cells treated with an sgRNA targeting *I.s. Ctr1A* are more resistant to treatment with CuCl_2_ than cells treated with a non-targeting (NT) sgRNA. (E) Small-scale pooled CuCl_2_ resistance assay. Cells were transfected with either of two *I.s. Ctr1A*-targeting sgRNAs or any of six NT sgRNAs, then cultured +/- 60 μM CuCl_2_ supplementation. After 48 days, *Ctr1A-*targeting sgRNAs were selectively enriched in the CuCl_2_-treated population. (F) Gene-level results of the genome-wide CuCl_2_ resistance screen (replicate 1 vs replicate 2). (G) Clustal Omega analysis of CTR1 family proteins: *Saccharomyces cerevisiae* (Sc) CTR1, CTR2, and CTR3; *Ixodes scapularis* (Is) Ctr1A and Cup2; *Drosophila melanogaster* (Dm) Ctr1A, Ctr1B, and Ctr1C; and *Homo sapiens* (Hs) SLC31A1 (also known as CTR1) and SLC31A2 (also known as CTR2). (H) Competition assay showing enrichment of sgRNAs targeting the indicated gene (*Ctr1A* or *Cup2*) or a non-targeting sgRNA (NT). Y-axis, fold enrichment of sgRNA-expressing cells in treated vs. untreated cell populations.

Based on conservation and structural analyses, we predicted that KO of *Ctr1A* would confer resistance to treatment of ISE18 cells with toxic levels of copper chloride (CuCl_2_). To test this, we first established levels of CuCl_2_ media supplementation that confer toxicity to *I. scapularis* ISE18 cells. We then tested the prediction that KO of *Ctr1A* would confer resistance to CuCl_2_ supplementation by challenging pools of cells receiving a non-targeting sgRNA (control) or a *Ctr1A*-targeting sgRNA with a range of levels of CuCl_2_ supplementation (**Fig. 3D**). As expected, introduction of a *Ctr1A*-targeting sgRNAs in RMCE+Cas9+ ISE18 DsRed cells resulted in resistance to CuCl_2_ as compared to the control (**Fig. 3D**). We next performed a small-scale pooled assay in which RMCE+Cas9+ ISE18 DsRed cells were first transfected with a given *Ctr1A*-targeting or non-targeting sgRNAs along with a PhiC31 integrase expression plasmid, then the transfected cell pools were combined and cultured in the presence or absence of CuCl_2_ supplementation (**Fig. 3E**). To assess enrichment of *Ctr1A*-targeting sgRNAs in the CuCl_2_-treated mini-pool, we analyzed the relative abundance of sgRNAs in the initial, treated, and untreated populations via PCR amplification and NGS. As expected, sgRNAs targeting *Ctr1A* were highly enriched in the CuCl_2_-treated cell pool but not in the untreated pool, increasing in proportion from ∼25% to ∼90% in the CuCl_2_-treated cell pool (**Fig. 3E**). This suggests that supplementation with CuCl_2_ provides an effective cell-based selection assay and that KO of at least one non-essential gene in the *I. scapularis* genome confers resistance to this treatment, making resistance to CuCl_2_ a promising proof-of-concept screen assay for testing of the genome-wide platform.

### Genome-wide screen identifies genes related to copper ion trafficking

To perform a CuCl_2_ selection screen at genome-wide scale, we subjected the KO cell library generated for the fitness screen to treatment with CuCl_2_, while maintaining in parallel control populations without copper supplementation (workflow in **Suppl. Fig. 3C**). After several weeks of continuous passage under untreated CuCl_2_-treated or control conditions, we harvested genomic DNA, amplified sgRNA cassettes by PCR, and quantified sgRNA abundance by NGS. As expected, gene-level analysis revealed reproducible enrichment of sgRNAs targeting *Ctr1A* in CuCl_2_-treated samples across both biological replicates (**Fig. 3F** and **Suppl. File 7)**. The most enriched gene in the CuCl_2_ resistance screen is annotated as “probable low affinity copper uptake protein 2” (LOC115329565; hereafter, Cup2) (**Fig. 3F**). Top-enriched genes also include orthologs of *Drosophila* genes with known or predicted roles in ion homeostasis or transport (i.e. orthologs of *D. melanogaster CG3392* and *nirvana 3*; **Suppl. File 7**). GSEA with PANGEA for *I. scapularis* identifies copper ion transport as the most-enriched molecular function or biological process (fold enrichment 127, P-value 6.157e-5; **Suppl. File 8**).

We were interested by the finding that KO of either of two genes encoding putative copper transporters, *Ctr1A* and *Cup2*, can make cells resistant to CuCl_2_ treatment. Results of InterProScan analysis suggest that like Ctr1A, Cup2 is also a Ctr copper transporter family protein (InterPro ID IPR007274) and that most of the protein is comprised of a single solute carrier family 31 copper transporter domain (Panther ID PTHR12483) ^44^. We next compared Cup2 protein sequences with *I.s.* Ctr1A and well-characterized Ctr family proteins from yeast, humans, and *Drosophila* using Clustal Omega ^49^. Among these proteins, *I.s.* Cup2 appears to be most closely related to human SLC31A2 (also known as CTR2; **Fig. 3G**). We then analyzed Cup2 structure with AlphaFold 3 ^46^. AlphaFold 3 results were inconsistent with formation of a Cup2 homotrimer (**Suppl. Fig. 5A**) but do suggest that Cup2 and Ctr1A can form a heterotrimeric complex. Indeed, structural predictions for 2 x Cup2 with 1 x Ctr1A and 1 x Cup2 with 2 x Ctr1A were consistent with a higher-confidence trimeric structure, and the addition of copper ion in the heterotrimer predictions further stabilized the complex structures (**Suppl. Fig. 5B,C**). Further, as we found for Ctr1A trimers (**Fig. 3A-C**), the Ctr1A-Cup2 heterotrimeric structures are predicted to bind copper in a central pore (**Suppl. Fig. 5B,C**) and show a preference for binding to copper over other positive ions, consistent with predicted roles as copper ion transporters (**Suppl. Fig. 5D,E**).

Finally, we sought to experimentally validate the finding that KO of *Cup2* results in copper CuCl_2_-resistant cells in a secondary cell assay. To do this, we performed a competition assay. We first transfected RMCE+ Cas9+ ISE18 DsRed cells with individual sgRNAs against *Ctr1A* or *Cup2,* or a non-targeting guide, in pLib025-iRFP720, along with a plasmid expressing PhiC31, resulting in a mix of sgRNA-expressing (iRFP720+) cells and non-transfected cells. Next, these mixed populations were treated with CuCl_2_ and cultured for several weeks. The ratios of no-sgRNA cells to with-sgRNA cells were compared for each cell population, before and after treatment. As expected for KO of a copper transporter, there was an enrichment of iRFP720 fluorescent protein-tagged cells expressing guides against either *Ctr1A* or *Cup2* but not the non-targeting guide after CuCl_2_ treatment (**Fig. 3H**). Altogether, these data provide experimental evidence consistent with the idea that *I. scapularis* Ctr1A and Cup2 are involved in copper transport and demonstrate that genome-wide KO screening using a CuCl_2_ selection assay successfully identified relevant genes.

### Screen for resistance to the bacterial metabolite Antimycin A

Ticks have adapted to long periods of fasting followed by rapid intake of nutrients in the form of a blood meal, placing unique metabolic demands on mitochondrial function ^50^. Furthermore, the mitochondrial electron transport chain plays a key role in pathogen infection through its regulation of cellular redox balance and metabolic state ^51^. With this in mind, we chose to screen for resistance to Antimycin A, a well-characterized bacterial metabolite that is an inhibitor of mitochondrial electron transport chain Complex III (reviews include ^52^) and for which a resistance screen dataset in human cells is available ^53^. To perform an antimycin A resistance screen, we treated genome-wide KO cell pool with Antimycin A while maintaining untreated controls in parallel, similar to the CuCl_2_ resistance screen (**Suppl. Fig. 3C**). Following multiple passages under selection, genomic DNA was harvested, sgRNA sequences were amplified by PCR, and relative abundance was quantified by NGS and subsequent data analyses.

The Antimycin A resistance screen data are provided as **Suppl. File 9** and a visualization of the data as **Suppl. Fig. 6**. As expected, the screen identified genes with well-established roles in mitochondrial function and cellular homeostasis, including LOC8025722 (*ND-B17.2*), which encodes a mitochondrial Complex I subunit and LOC8038256 (*Tmem186*), which encodes an ortholog of its assembly factor; LOC115324469, which encodes an ortholog of the Complex V component F1F0 ATP synthase; and genes that encode orthologs of proteins with roles in central metabolic pathways, including a bifunctional enzyme in CoA biosynthesis (LOC8025601, *Ppat–Dpck*) and a long-chain fatty acid CoA ligase (LOC120843733). We also compared the *I. scapularis* Antimycin A screen results with genes identified in a similar screen in human cells ^53^, and found that one *I. scapularis* gene, LOC8041162 (*akirin-2*), is related to a gene enriched in the human cell screen, *AKIRIN1*. These data further support the idea that resistance screens in *I. scapularis* cells are feasible and yield meaningful results.

### Genome-wide screen for resistance to the *Metarhizium* fungal toxin Destruxin A (DA) implicates the glycophosphatidylinositol (GPI) anchor synthesis pathway

Genome-wide screening offers a powerful approach to advance the development of tick control strategies. *Metarhizium* fungi hold promise for arthropod control, including tick control ^54–56^. These fungi secrete cyclic depsipeptides, including Destruxin A (DA), that have cytotoxic and other effects on cells ^28,57^. DA is known to affect calcium levels in insect cells ^58^. We reasoned that application of our pooled CRISPR screening approach to identify genes required for DA toxicity could identify genes required for entry of the toxin and/or mechanisms of DA cytotoxicity. In an *in vivo* assay, we found that immersion of *I. scapularis* nymphs in a solution containing DA results in reduced survival in DA-treated but not vehicle control-treated nymphs (**Fig. 4A**), suggesting that a DA resistance screen can yield *in vivo* relevant results.

**Figure 4.**
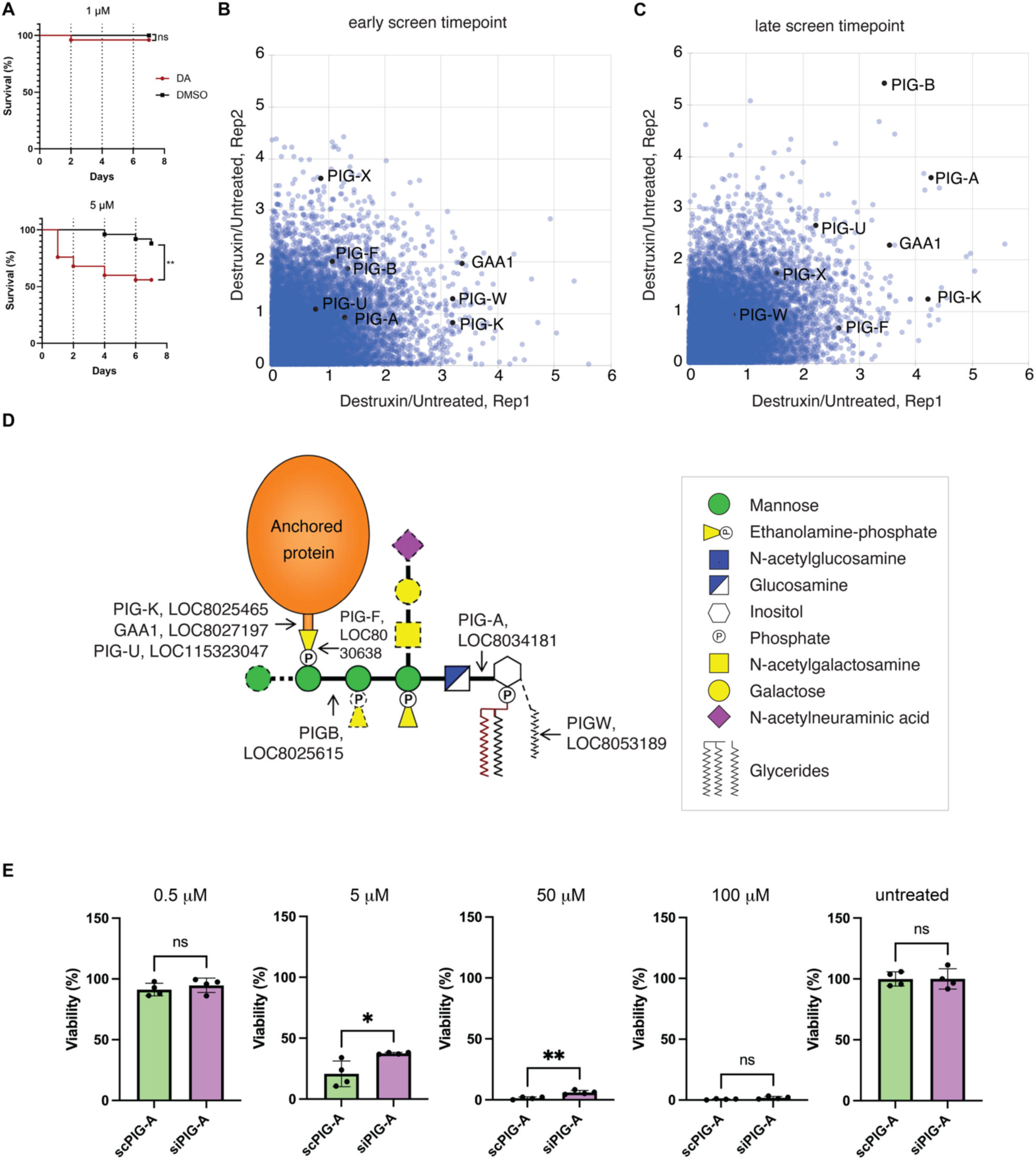
(DA-screen): Genome-wide screen for resistance to the fungal toxin Destruxin A (DA) identifies genes encoding glycosylphosphatidylinositol (GPI) anchor synthesis pathway components. (A) Immersion of *I. scapularis* nymphs into a solution containing DA reduces survival as compared with a vehicle control. Top, 1 uM dose of DA. Bottom, 5 uM. Red, immersion in a solution treated with DA (stock solution dissolved in DMSO). Black, vehicle control (equal volume of DMSO). (B-C) Plots of negative log of the enrichment scores in each replicate of the DA resistance screen at an early (B) and a late (C) timepoint in the assay. Genes are indicated using human gene symbols; for gene mapping see

To perform the DA screen, cells were exposed to the toxin across several passages (see **Methods** and **Suppl. Fig. 3C**), and sgRNA abundance was quantified by NGS at both early and late timepoints in the screen (**Fig. 4B,C** and **Suppl. File 10**). Consistent with prior observations that DA exposure induces calcium influx and suggesting that modulation of calcium homeostasis can confer resistance ^58^, we found that a predicted calcium pump (LOC8025060, *ATP8B*) is enriched in DA-treated cells (**Suppl. File 10**). The screen data also reveals robust and reproducible enrichment of multiple genes involved in GPI-anchor biosynthesis in DA-treated cell KO populations (**Fig. 4B-D** and **Suppl. File 10**). Using the *I. scapularis* version of PANGEA for GSEA (minimum enrichment score ≥ 2), the top enriched category is “attachment of GPI anchor to protein” (150-fold enrichment; P-value 7.15964E-05; **Suppl. File 11**). A summary of human GPI anchor biosynthesis genes, their predicted orthologs in *I. scapularis*, and scores for the orthologs of these genes in the *I. scapularis* cell screen is provided in **Table 2** and an annotated diagram of the canonical pathway is shown in **Fig. 4D**.

**Table 2:** Human GPI anchor biosynthesis pathway genes and their orthologs in *I. scapularis.* (D) Summary illustration of the canonical GPI anchor synthesis pathway with human gene names and corresponding *I. scapularis* ortholog NCBI Gene IDs indicated. Adapted from Liu et al. A knockout cell library of GPI biosynthetic genes for functional studies of GPI-anchored proteins, 2021 Communications Biology35 (CC-BY-4.0). (E) RNAi knockdown of *I. scapularis PIG-A* in ISE6 cells in the presence or absence of treatment with DA at the indicated concentrations, comparing results of MTT assay with a scrambled control RNA (scPIG-A) vs. a *PIG-A*-targeting short interfering RNA (siPIG-A). Knockdown of *PIG-A* confers statistically significant resistance to treatment with DA. *, P-value 0.0489. **, P-value 0.0032.

| *Homo sapiens* | *Drosophila melanogaster* | *Ixodes scapularis* | DA screen data (Rep 1)<sup>1</sup> | DA screen data (Rep 2)<sup>1</sup> |
| --- | --- | --- | --- | --- |
| **PIG-A** | **PIG-A** | **LOC8034181** | **4.28** | **3.6** |
| **PIG-B** | **PIG-B** | **LOC8025615** | **3.45** | **5.42** |
| PIG-C | PIG-C | LOC115319264 | 0.54 | 0.47 |
| **PIG-F** | **PIG-F** | LOC8030638 | **2.6** | 0.64 |
| PIG-G | PIG-G | LOC115310841 | 0.8 | 1.06 |
| PIG-H | PIG-H | LOC8044011 | 2.12 | 1.49 |
| **PIG-K** | **PIG-K** | **LOC8025465** | **4.23** | 1.23 |
| PIG-L | PIG-L | LOC8032204 | 0.81 | 0.68 |
| PIG-M | PIG-M | LOC120851149 | 2.12 | 1.44 |
| PIG-N | PIG-N | LOC8028039 | 1.59 | 0.4 |
| PIG-O | PIG-O | LOC8028655 | 1.02 | 0.52 |
| PIG-P | PIG-P | LOC8024414 | 1.07 | 1.55 |
| PIG-Q | PIG-Q | LOC8033187 | 0.28 | 0.64 |
| PIG-S | PIG-S | LOC120846754 | 2.03 | 0.52 |
| PIG-T | PIG-T | LOC8032324 | 1.05 | 1.19 |
| **PIG-U** | **PIG-U** | LOC115323047,<br>LOC115330552 | 2.25, 2.18 | **2.67**, 1.67 |
| PIG-V | PIG-V | LOC8027599 | 0.56 | 0.51 |
| **PIG-X** | **PIG-X** | LOC115328618 | 1.49 | 1.68 |
| PIG-Y | CG46311 |  |  |  |
| PIG-Z | PIG-Z | LOC8040615 | 0.31 | 0.82 |
| **PIGW** | **PIG-Wa,b** | LOC8053189 | 0.72 | 0.9 |
| **GAA1** | **GAA1** | LOC8027197 | **3.55** | 2.28 |
| B3GALT4 | CG3038 | LOC8040327 | 1.19 | 1.33 |
| DPM1 | Dpm1 | LOC8025171 | 0.02 | 0 |
| DPM2 | Dpm2 |  |  |  |
| DPM3 | Dpm3 | LOC8026541 | 0.17 | 1.09 |
| PGAP1 | PGAP1 | LOC8033123 | 1.1 | 0.05 |
| PGAP2 | PGAP2 | LOC115320393,<br>LOC115320391 | 1.94, 0.15 | 1.60, 0.47 |
| PGAP3 | PGAP3 | LOC8034135 | 0.8 | 0.89 |
| PGAP4 |  |  |  |  |
| PGAP5 |  |  |  |  |

<sup>1</sup> Negative log enrichment scores in each replicate of the *I. scapularis* DA resistance screen.
**Bold italics** indicates that the *I. scapularis* gene meets our criteria for a hit in the DA resistance screen.

To confirm the relevance of the GPI anchor biosynthesis to DA toxicity, we assessed if RNAi knockdown of *PIG-A*, which encodes the ortholog of a protein that acts early in the GPI synthesis pathway (**Fig. 4D**), makes cells resistant to treatment with DA. To do this, we treated ISE6 cells with siRNAs targeting *PIG-A* and measured viability following DA treatment. Relative to control siRNA-treated cells, *PIG-A* knockdown increased viability across multiple DA concentrations (**Fig. 4E** and see **Suppl. Fig. 7**), supporting a role for PIG-A and the GPI anchor biosynthesis pathway in mediating sensitivity of *I. scapularis* cells to DA. These results suggest that GPI-anchor biosynthesis genes are determinants of DA susceptibility and demonstrate the utility of our platform for dissecting host-toxin interactions in *I. scapularis* cells.

## Discussion

Application of ‘omics technologies such as genome and transcriptome sequencing allow for relatively rapid and precise annotation of genes encoded by non-model species, including non-model arthropods such as ticks. However, only a subset of genes can be assigned putative function based on similarity to genes of known function in other species and even for conserved genes, experimental data is lacking. Moreover, although reverse genetic approaches are useful for testing hypotheses regarding gene function, methods for unbiased screening are needed in order to understand differences among species, uncover species-specific or tick-specific gene activities, and functionally annotate non-conserved genes. Indeed, uncovering the unique biology of non-model organisms is a key goal for organisms such as ticks, as understanding these differences can aid development of strategies that mitigate tick-borne diseases through tick control. In this work, we showed that CRISPR-Cas9 KO cell screening can be successfully extended to cell lines derived from the tick *I. scapularis.* The engineered cells, plasmid vectors, and genome-wide sgRNA library we report here comprise a robust screen platform useful for study of a variety of cell biological topics. For many genes, the screen datasets we report provide the first experimental evidence of function obtained in an *I. scapularis* system.

Identification of fitness genes informs interpretation of subsequent screens by providing a list of genes unlikely to be identified because they are required for cell growth and/or survival. In addition, fitness genes might be of particular interest for disease vectors such as *I. scapularis,* as these genes serve as potential targets for development of small molecule, peptide, or other approaches to vector control. Conserved genes that scored as fitness genes in our screen in *I. scapularis* ISE18 cells but not in *D. melanogaster* S2R+ cells might point to unique vulnerabilities in ticks; however, this set of genes is also likely to reflect the different cell types investigated. ISE18 cells are similar in morphology and transcriptome to ISE6 cells, which are neuronal-like cells ^59^. Genes that score as fitness genes in ISE18 cells but not in hemocyte-like *Drosophila* S2R+ cells include *I. scapularis* orthologs of *Drosophila* neuronal genes such as *elav*. Follow-up studies will be needed in order to separate cell type-specific vs species-specific dependencies, validate conserved putative fitness genes *in vivo* in *I. scapularis*, and prioritize targets for vector control.

Another category of fitness genes of particular interest are the subset of fitness genes that are poorly conserved in other species (for comparison with *Drosophila* and human genes, see **Suppl. File 1**). Among these, we suggest that LOC120848847 (VectorBase ID ISCP_029989) and LOC115322987 (ISCP_023786) might be prioritized for follow-up. These genes meet our criteria for fitness genes (cutoff of FDR 0.1) and are expressed in ISE18 cells, consistent with what is predicted for true hits. These genes are up-regulated in the female midgut following post-engorgement, consistent with potential roles in cell growth and/or proliferation (data reported by ^60^ and mined at VectorBase ^61^). Both genes encode relatively short open reading frames (ISCP_029989, 101 amino acids and ISCP_023786, 148 amino acids) with no identified functional domains and no readily detectable orthologs outside of the *Ixodes* genus.

Selection-based assays (resistance screens) have been particularly informative in *Drosophila* and mosquito cell studies ^24–26,62^. To establish proof-of-concept for selection-based assays, we screened for KO cells that are resistant to CuCl_2_ or Antimycin A. As expected, the CuCl_2_ screen identified *I. scapularis Ctr1A*, a gene we had established in a small-scale test could confer resistance to CuCl_2_, and another Ctr1 family transporter, *I. scapularis* Cup2. Similar to the CuCl_2_ screen, the screen with Antimycin A resulted in identification of expected hits, further indicating that resistance assays are feasible in this system.

Our CuCl_2_ screen revealed that KO of either *Ctr1A* or *Cup2* can confer resistance to the treatment, suggesting that these genes have non-redundant functions. The results of AlphaFold modeling suggest that Ctr1A and Cup2 are capable of forming heterotrimers that show preference for binding of a copper ion in a central pore. Thus, one plausible hypothesis to explain the non-redundant function of these genes in our assay is that Ctr1A and Cup2 physically interact. However, we cannot rule out alternative explanations, such as differences in subcellular localization of the two proteins, and we note that in mammals, CTR family proteins have been shown to exert regulatory control on one another ^63,64^. Relevant to tick control, the experimental finding that KO of either *Ctr1A* or *Cup2* can confer resistance to treatment with cytotoxic levels of copper might provide insights into potential routes for resistance of whole ticks to copper and other metal ion nanoparticle-based tick control approaches. Ion nanoparticle-based control has begun to be explored as a method for tick control ^65,66^ and is of particular interest due to new availability of “green” methods for nanoparticle synthesis (e.g., see ^67,68^).

Exposure of ticks to pathogenic fungi and/or the biotoxins they encode provides another potential control strategy, and understanding toxin entry can help inform potential applications. Our selection-based assay with the *Metarhizium* fungal toxin DA did not yield an obvious candidate cell-surface receptor; however, components of the GPI synthesis pathway are clearly enriched in the screen, suggesting that one or more GPI-anchored protein acts as a receptor of DA or is otherwise required for sensitivity to the toxin. This is consistent with what has been found for some other microbial toxins. For example, GPI anchors are a common feature of receptors of the bacterial toxin aerolysin ^69^ and GPI anchor synthesis genes were identified in a gene trap-based genetic screen for aerolysin resistance in haploid mammalian cells ^70^. Moreover, a retroviral mutagenesis screen identified GPI synthesis pathway genes as required for toxicity of *Clostridium* alpha toxin to Chinese hamster ovary (CHO) cells ^71^, and a CRISPR screen for resistance to β-hemolysin/cytolysin (BHC), a pore-forming toxin produced by a *Streptococcus* bacterial species, identified GPI synthesis pathway genes as required for BHC toxicity to HeLa cells ^72^. Detection of multiple critical components of the GPI synthesis pathway in the *I. scapularis* screen highlights the sensitivity of our platform.

Although our datasets clearly show that the *I. scapularis* CRISPR cell screening platform is effective, there remains room for improvement. Regarding sgRNA design, the results we obtained suggest that annotations are sufficient for discovery of many relevant factors. Nevertheless, availability of improved gene models would improve sensitivity of the screens, given that incomplete or incorrect gene models can confound sgRNA design and gene-level analysis of screen data. In addition, as we accumulate additional data, we will be able to apply a machine learning approach to discover rules for sgRNA design that might be different from rules learned based on studies in cells from other species. In the future, we can also use the ISE18 and ISE6 cell genome sequence data to exclude sgRNAs with SNPs in cells as compared with the reference genome prior to sgRNA synthesis. In addition, developing a robust single-cell cloning protocol would likely be useful both for screening and for other applications, and improving transfection efficiency would enhance screen quality and reduce costs. Moreover, expanding the platform to additional *I. scapularis* cell lines with distinct transcriptional profiles and pathogen responses would facilitate separation of general fitness genes from those that are cell type-specific, as well as a broader diversity of screen assays. Regarding assay design, in the current version of the platform, we relied heavily on FACS and used multiple fluorescent channels, which might complicate development of assays using fluorescent-tagged microbes or transcriptional reporters; to address this in part, we engineered a second, iRFP670 version of the platform, freeing up the DsRed channel. The system could also be further improved by development of reliable protocols for antibiotics-based enrichment. Furthermore, development of a platform for over-expression screening would circumvent some limitations of screening with only one or a few cell types.

Altogether, we successfully developed a robust CRISPR KO screening platform for *I. scapularis* cells, opening the doors to unbiased forward-genetic screening in cells from a relatively under-studied but highly human health-relevant species. We anticipate that the platform will be used in the future for increasingly specific and relevant assays, and that further refinement of the platform will contribute to increased precision and sensitivity of future *I. scapularis* cell screens.

## Methods

### Cell culture

The *Ixodes scapularis* cell lines ISE18 and ISE6 ^8^ were kind gifts from Ulrike Munderloh (University of Minnesota). All cell lines were maintained at 34°C in freshly prepared L15C300 tick medium ^8,73^ with 10% heat inactivated fetal bovine serum (Gibco), DIFCO Tryptose Phosphate Broth, Bacto Buffered Medium (BD), and Lipoprotein-cholesterol concentrate (MP Biomedicals).

### Pol II activity assays

Cells were seeded at a density of 1 × 10^!^cells per well in 1 mL of complete culture medium in 12-well plates and allowed to adhere. Cells were co-transfected with 0.5 µg of a firefly luciferase reporter plasmid driven by the promoter of interest and 0.5 µg of a constitutive CAG-driven *Renilla* luciferase plasmid as a normalization control. Plasmid DNA was diluted in 50 µL Opti-MEM (Thermo Fisher Scientific) and combined with 2 µL P3000 reagent. In parallel, 3 µL Lipofectamine 3000 (Thermo Fisher Scientific) was diluted in 50 µL Opti-MEM. The two solutions were mixed and incubated for 10 minutes at room temperature before being added dropwise to cells. Three days post transfection a dual luciferase assay was performed using the Dual-Glo Luciferase Assay System (Promega) according to the manufacturer’s instructions, with CAG promoter-driven Renilla luciferase as the normalization control. Firefly and *Renilla* luminescence were measured using a plate reader (Molecular Devices Spectramax Paradigm). For each well, firefly luciferase activity was normalized to *Renilla* luciferase activity to control for transfection efficiency and cell number. Relative promoter activity was calculated as the ratio of firefly to *Renilla* luminescence.

### Codon optimization of open reading frames (ORFs) for expression in *I. scapularis*

Genome-scale codon preference in *I. scapularis* was estimated based on the protein-coding gene annotation (GCF_016920785.2_ASM1692078v2_cds_from_genomic.fna.gz), downloaded from the NCBI genome FTP site, by counting the frequency of each codon used, then setting a cut-off for rare codons as <1% frequency. Next, we grouped the results based on amino acid and for each amino acid, we separated the codons into two classes, rare codons and non-rare codons, then built an annotation file. An in-house program was written to process ORFs of interest by evaluating each codon based on the annotation file and then replacing each rare codon with a randomly selected non-rare codon for the same amino acid. The pipeline was applied to the PhiC31 open reading frame and a codon-optimized version of the ORF was then commercially synthesized (Genscript).

### Cell engineering

To generate RMCE+ Cas9+ ISE18 and ISE6 tick cell lines, we first constructed a plasmid derived from pT/CAGGS-DsR//CMV-SB, which carries both Sleeping Beauty (SB) transposon and transposase functions ^74^. The region inside the transposon inverted repeats was modified to include an RMCE cassette consisting of attP sites flanking a CAG promoter-driven expression unit with open reading frames (ORFs) for DsRed and Zeocin resistance (SBFinal_attP_CAG_DsRed_Zeo_attP) separated by a T2A sequence. An iRFP670 version was also constructed by modifying the pT2/SVNeo cassette (Addgene ID 26553) to contain an ORF for iRFP-670 and blasticidin resistance separated by T2A (pT2SV-attP-Cag-iRFP-Blast-attP). Plasmids were validated by whole-plasmid sequencing (Plasmidsaurus) and transfected into ISE6 or ISE18 cells using Lipofectamine 3000 (Thermo Fisher Scientific). The cassette pT2SV-attP-Cag-iRFP-Blast-attP was co-transfected with a second plasmid expressing the SB transposase pCMV/SB10 (Addgene ID 24551).

Four days post-transfection, cells were enriched by fluorescence-activated cell sorting (FACS) using a MoFlo Astrios EQ cell sorter (Beckman Coulter) with a 100 µm nozzle at the Harvard Medical School (HMS) Immunology Flow Cytometry Core Facility. DsRed or iRFP-positive cells were collected based on broad gating to obtain 4.5 million cells to fill a T25 flask. The sorted population was expanded to 12 T25 flasks over several weeks and enriched again. Three additional rounds of DsRed enrichment were then performed (for a total of four rounds of enrichment). For the first, second, and third rounds, DsRed-negative cells were added after sorting to maintain a total of 4.5 million cells per flask. In the final round, we were able to fully enrich the population, sorting 4.5 million DsRed positive cells to fill a T25 flask.

Next, to generate a Cas9-expressing version of the RMCE (DsRed or iRFP670) cell population, each of six T25 flasks were seeded with 9 million RMCE+ ISE6 or ISE18 cells and transfected with a custom SB vector encoding a CAG-driven Cas9-2A-Neo-T2A-TagBFP cassette (VectorBuilder), along with a source of SB transposase, pCMV/SB10 (Addgene ID 24551). Four days post-transfection, cells were enriched by FACS for dual DsRed+ and TagBFP+ expression using the same sorter and strategy. Gating was again set to recover the top 4.5 million fluorescent cells to fill a T25 flask. The resulting polyclonal, dual-positive population was expanded and maintained for use in CRISPR screen assays.

### Confirmation of RMCE in ISE18 cells

To test if introduction of PhiC31 and an appropriate cassette can mediate RMCE in RMCE+ ISE18 cells, we tested whether DsRed could be exchanged for GFP in the presence of a donor cassette, pLib-CAG-GFP-P2A-Puromycin flanked by attB sites (pLib-GFP), and CAG-PhiC31 integrase. RMCE+ Cas9+ ISE18 DsRed cells were plated at 70–80% confluency in two T25 flasks. One flask was transfected with pLib-GFP and a CAG-driven PhiC31 integrase expression plasmid and a second flask was transfected with pLib-GFP and an empty vector as a control. Transfections were performed using Lipofectamine 3000 (Thermo Fisher Scientific) according to the manufacturer’s instructions. For each flask, a P3000 mix was prepared by combining 250 µL Opti-MEM, 20 µL P3000 reagent, and 8 µg total DNA (4 µg of pLib-GFP donor and 4 µg of PhiC31 integrase or control plasmid). Separately, a Lipofectamine mix was prepared by combining 250 µL Opti-MEM and 20 µL Lipofectamine 3000. The Lipofectamine mix was added to the DNA/P3000 mix, incubated for 10–15 minutes at room temperature, then the complexes were added to cells. The culture media was replaced the following day. On days 4 and 36 post-transfection, cells were analyzed by flow cytometry using a BD LSR-II at HMS Immunology Flow Cytometry Core Facility.

Blank cells and single-color controls (DsRed-only, GFP-only, and TagBFP-only) were used to set gates and compensation. Then, gating was performed on sub-population of experimental or control cells that were TagBFP+ cells. We used gain of GFP expression in TagBFP+ cells as an indicator of successful RMCE. An empty vector control condition was included to account for background recombination and transient plasmid expression.

### Transcriptomics analysis

To prepare cell samples, the cell lines IDE2, ISE6, and ISE18, as well as an enriched pool of RMCE+ ISE18 DsRed cells, were seeded on T25 flasks and incubated at 34°C for 4 days to a confluency of ∼60%. On day 4, cells were washed and counted on Countess to prepare samples for RNA sequencing. Four replicates of each sample were collected, with two replicates derived from each flask consisting of 5.5 × 10^6^ cells total. Next, to prepare RNA, Eppendorf tubes containing the combined cell suspensions were centrifuged at 200 rcf for 4 minutes to pellet the cells. The supernatant was carefully aspirated, ensuring the cell pellet remained intact in the tube. To each tube, 600 µL of TRIzol reagent was added, and the contents were gently pipetted up and down to resuspend and break up the pellet. The tubes were then placed in a −80°C freezer for storage. On Day 5, the frozen samples were retrieved from −80°C for RNA extraction. RNA isolation was performed using Direct-zol RNA Miniprep Kit (Zymo Research).

RNA was then poly-A selected, quality analyzed, and subjected to NGS at the Biopolymers Facility at Harvard Medical School (RRID SCR_007175). For all samples, the results of electrophoretic analysis profiles and quantitative PCR (qPCR) were consistent with high-quality RNA. KAPA mRNA HyperPrep Kits were used to poly-A select mRNA libraries. As described for the public data deposition (NCBI GEO Accession GSE269712), NGS was performed using an Illumina NovaSeq 6000 sequencing system with an SP flow cell, and reads were mapped to the *I. scapularis r*eference genome (GCF_016920785.2) using STAR version 2.7.9a and gene count extraction using FeatureCounts version 2.0.1. Normalization and conversion to FPKM values were performed using DEseq2 version 1.40.2. To compare among cell lines, we compiled a list of 10,645 genes detected as expressed in any of the samples. Next, the FPKM values for these genes were used to determine the Pearson correlation coefficient between samples. The correlation coefficients for all pairs of samples were then plotted as a heatmap.

### ISE18 cell genome sequencing and SNP analysis

Wild-type ISE18 were pelleted at 200 × g and frozen and shipped on dry ice to Novogene. DNA-Seq was performed by Novogene using a single lane on a Novoseq X Plus. Raw data processing and the SNP calling were done by Novogene using the bcftools v1.16 pipeline. We defined differences between these data and the NCBI reference genome for *I. scapularis* as SNPs if the difference was supported by at least ten reads. The genome locations of the selected SNPs were compared with the sgRNA target locations on the genome, including the PAM. An sgRNA was removed from the library we define as the “version 1.1” library if any of the selected SNPs fall in the target location.

### Pol III promoter identification and activity assays in ISE18 cells

To identify candidate U6 promoters in ticks, the *D. melanogaster* U6:2 (CR32867) snRNA sequence was used as a query in BLAST searches against *Ixodes scapularis* genome assemblies at Vectorbase (IscaW1.7 and IscaI1.0) ^61^. Orthologous loci were aligned using Clustal Omega ^49^, and regions including the snRNA sequence with 300 bp upstream and 50 bp downstream were examined for canonical Pol III promoter elements. Four candidate promoters were selected based on the presence of predicted promoter features, and regions spanning 220 bp upstream and 50 bp downstream of the snRNA were synthesized in gBLOCKs (IDT) and cloned into a backbone library vector, i.e., pLib6.6b (Addgene ID 176666) modified to express GFP under control of a CAG promoter. The pol III candidate promoter sequences are provided in **Suppl. File 12 (Materials)**. To evaluate activity, RMCE+ Cas9+ ISE18 DsRed cells were transfected with plasmids expressing sgRNAs targeting DsRed driven with each candidate promoter or a *D. melanogaster* pol III promoter. Specifically, cells were plated at 70–80% confluency in T25 flasks and transfected with one of the following: pLib-GFP/NT (non-targeting control), pLib025-GFP/DsRed, pLib087-GFP/DsRed, pLib518-GFP/DsRed, pLib949-GFP/DsRed (encoding tick promoters), or pLibDM-GFP/DsRed (*D. melanogaster* U63). Transfections were performed as described for the RMCE assay. Briefly, for each pLib construct, 8 µg of the construct was transfected into a separate T25 flask of RMCE+ Cas9+ ISE18 DsRed cells using Lipofectamine 3000 (Thermo Fisher Scientific). Media was replaced the following day. Cells were maintained by splitting or media change as they approached confluency, typically once per week. Flow cytometry analysis was performed on days 10 and 20 post-transfection using a BD LSR-II analyzer. Gating was performed on TagBFP+ and GFP+ cells (to select for cells expressing Cas9 and the sgRNA plasmid), and within this population, DsRed median fluorescence intensity was measured as a readout of KO efficiency.

### AlphaFold 3 structural predictions for *I. scapularis* CTR family proteins

Protein complex predictions were performed using AlphaFold 3 ^46^. To determine the optimal oligomeric state of *Ixodes scapularis* Ctr1A, we predicted complexes containing 2, 3, 4, and 5 subunits of the Ctr1A protein sequence. For each stoichiometry, five models were generated using the AlphaFold 3 server default settings (random seed). Predicted complexes were evaluated using two complementary metrics: ipTM (interface predicted Template Modeling score; ^47^) and iLIS (integrated Local Interaction Score), computed using the AFM-LIS jupyter notebook pipeline (https://github.com/flyark/AFM-LIS) ^75^ and scores were averaged across the five models. The same approach was used to assess models for Cup2 trimers or Ctr1A-Cup2 heterotrimers.

To assess ion binding selectivity, the Ctr1A homotrimeric complex was predicted in the presence of each ion species available in the AlphaFold 3 server (Cu²⁺, Co²⁺, Fe²⁺, K⁺, Na⁺, Mn²⁺, Cl⁻, Ca²⁺, Zn²⁺, and Mg²⁺), with five models generated per ion. iLIS and ipTM scores were computed for each prediction to compare ion binding propensity. Additionally, predictions of the Ctr1A trimer with and without Cu²⁺ were compared by examining per-residue pLDDT (predicted Local Distance Difference Test) scores to assess the effect of copper binding on predicted structural confidence. All structural visualizations were generated using UCSF ChimeraX ^76^. The same approach was used to assess predicted ion binding for Ct1A, Cup2 heteromeric complexes.

### Assay of sensitivity of Ctr1A KO cell pools to CuCl_2_

We used total ATP levels as a proxy for cell number and viability to assess sensitivity to CuCl₂. To do this, we created 96-well plates with a dilution series of CuCl₂ (2-fold dilution series), then added 70 µL of cells suspended in media. Enriched pools of cells that received either a non-targeting sgRNA or a *Ctr1A*-targeting sgRNA (i.e. pool of CTR1A KO cells; 30,000 cells per well) were added to 96-well plates, bringing the final volume per well to 140 µL. After a 7-day incubation, total ATP levels were assayed using Promega CellTiter-Glo (CTG). To do this, 90 microliters of CTG was added to each well, the plate was shaken for 5 min at 90 rpm, followed by a 10-min incubation at room temperature. Luminescence was then measured on a plate reader (Molecular Devices Spectramax Paradigm).

### Small-scale pooled assay for CuCl₂ resistance

Approximately 9 × 10⁶ RMCE+ Cas9+ ISE18 DsRed cells were seeded into a T25 flask in 5 mL of tick medium. Cells were transfected with a plasmid mixture containing equimolar amounts of CAG-PhiC31 and the sgRNA donor plasmid library pLib025-GFP, which encoded either one of two sgRNAs targeting *Ctr1A* or one of six non-targeting control sgRNAs, using Lipofectamine 3000. Pools of GFP positive cells (i.e. cells with integrated sgRNAs) were enriched by FACS two times over eight weeks. After enrichment, cells were counted using a Countess cell counter (Thermo Fisher) and then combined in equal proportions. An aliquot of this cell mixture was collected, pelleted at 200 × g, and stored at −80°C; the remaining cell mixture was divided evenly into two T25 flasks. One flask was treated with 60 µM CuCl₂ and the other was maintained in normal media. Cells were cultured for 48 days with weekly passaging and regular media changes. After 48 days, cells subject to CuCl₂ and control conditions were pelleted at 200 × g and genomic DNA extracted. Next, integrated sgRNAs were amplified by PCR from each sample. PCR products were then purified and submitted for NGS at the MGH CCIB DNA Core. Finally, the sgRNA representation in the initial, final untreated population, and final 60 µM CuCl₂-treated populations were quantified and compared.

### Genome-wide sgRNA library design

Genome wide knockout sgRNAs targeting the CDS region of the protein-coding genes from *Ixodes scapularis* were designed using the previously described pipeline ^19^ based on the reference genome annotation at NCBI RefSeq (GCF_016920785.2) and made available at our GuideXpress online resource ^20^. Up to 7 sgRNAs were selected based on the specificity and efficiency, considering the minimal OTE (off-target effect) score and maximum ML (machine learning efficiency) score. Any sgRNAs with *Bbs*I sites were removed. We also removed any sgRNAs targeting genes that failed to be detected in all three cell lines for which we did RNA-seq (FPKM cutoff 0.1). SNP calling results supported by at least 10 reads coverage based on the ISE18 DNA-seq data and for which SNP(s) were detected in the target region were removed virtually (v1.1 library design) for use in data analysis.

### Genome-wide sgRNA library cloning

Custom oligonucleotide pools encoding sgRNAs were synthesized as single-stranded DNA chips (Agilent). Each chip was dissolved in 200 µL of nuclease-free water, and half of the material (100 µL) was used directly as template for library amplification, with the remaining 100 µL stored as backup. Amplification of the library was performed by PCR using Phusion polymerase (New England Biolabs). Briefly, a 200 µL reaction mixture containing 40 µL water, 40 µL reaction buffer, 16 µL dNTP mix (2.5 mM each; Takara, Cat#4030), 1 µL of each primer (100 µM forward and reverse), 2 µL Phusion Hot Start Flex polymerase (New England Biolabs), and 100 µL dissolved chip DNA was prepared. To minimize amplification bias, the reaction mixture was divided into ten 20 µL PCRs. PCR was carried out with an initial denaturation at 95°C for 2 min, followed by 17 cycles of 95°C for 15 s, 59°C for 15 s, and 72°C for 30 s, with a final extension at 72°C for 10 min. Amplified material from the 10 reactions was pooled and resolved by agarose gel electrophoresis. The desired library fragment was purified from the gel using QIAEX II resin (Qiagen) and eluted in 20 µL nuclease-free water. Purified amplicons were cloned into the library vector backbone using NEBuilder HiFi DNA Assembly Master Mix (New England Biolabs). Vector DNA and purified insert were combined and run at 50°C for 1 h. Assembly products were subsequently transformed into high-efficiency electrocompetent cells 10GF’ ELITE Electrocompetent Cells (Lucigen) and a small portion was plated onto 10 cm LB plates containing carbenicillin and grown overnight at 30°C., ensuring recovery of at least 10–50× the theoretical library diversity. The rest was added to LB broth and grown for 12-16 hours at 30°C. Plasmid DNA was prepared from cultures using standard Qiagen midi-prep protocols, yielding the final sgRNA library (pLib025-GFP/GW).

### Generation of a genome-wide *I. scapularis* KO cell library and fitness assay

We generated a genome-wide KO cell library two independent times with minor changes. For both, approximately 9 million RMCE+ Cas9+ ISE18 DsRed cells were seeded in a T25 flask in 5 mL of tick media. Cells were then transfected with a plasmid mixture containing equimolar amounts of CAG-PhiC31 and the sgRNA donor plasmid library (pLib025-GFP/GW) using lipofectamine 3000. In the first set of screens, “genome-wide 1” (GW1), we began with 80 transfected flasks and carried forward 20 T25 flasks for expansion and characterization; in the second, “genome-wide 2” (GW2), we transfected 200 T25 flasks. A codon-optimized version of PhiC31 was used in GW2. For both, 4 days after transfection, cells were sorted by FACS at the HMS Immunology Flow Cytometry Core Facility for Cas9/TagBFP+ and pLib-GFP+. Cells were expanded (as free plasmid was presumably lost), then enriched a second time by FACS at the same facility. Cells were then grown in media supplemented with 0.45 ug/mL puromycin. Fitness assays were performed 59 days (GW1) or 56 days (GW2) after transfection. The GW1 or GW2 KO cell pools were then kept in culture and used for selection assays (below).

### Selection (resistance) assays with the genome-wide *I. scapularis* KO cell library

The GW1 KO cell pool was used for CuCl_2_ and DA screens, and the GW2 KO cell pool was used for CuCl_2_ and antimycin screens. For selection assays, cells were transferred to a flask containing 0.045 ug/mL puromycin and (i) CuCl_2_ dissolved in media to a final concentration of 60 uM at day 59 (GW1) or day 56 (GW2) post transfection, (ii) DA added to the media to at indicated concentrations (see below) at day 67 post transfection (GW1 only), or (iii) treated with 2.5 nM Antimycin A at 84 days post transfection (GW2 only). For the CuCl_2_ resistance assay, cells were continually passaged in the selective medium for ∼8 weeks (GW1) or ∼11 weeks (GW2); the timing was adjusted based on recovery of cell population following the initial treatment with CuCl_2_. For DA (GW1 only), cells were treated with 800 nM DA, allowed to recover in standard media for 6 wks, treated with 400 nM DA for 2 wks, then treated again with 800 nM DA for 3 wks. For antimycin a (GW2 only), cells were treated with Antimycin A-containing media (2.5 nM) for 44 days, allowed to recover in standard media for 1 week, returned to selective media for 10 days, allowed to recover again in standard media for 1 week, and returned to selective media for an additional 4 weeks. In all cases, re-seeding density was maintained at ∼1000 cells/sgRNA in 15 T25 flasks (GW1) or 3 t-175 flasks (GW2). In all cases, cells were resuspended or split into fresh media every 5-6 days, and matched untreated controls were maintained in standard media on the same schedule. At the start-point and endpoints for each screen, a sample with >1000 cells/sgRNA was pelleted and frozen at −80°C.

### Sample preparation, NGS, and analysis for genome-wide screens

For each cell pellet and the sgRNA plasmid libraries used in the screens (GW1 or GW2), genomic DNA was extracted using Quick-gDNA MiniPrep kits (Zymo). Next, genomic DNA was subjected to 2-step PCR to introduce in-line barcodes, a variable fingerprint, and Illumina sequencing primer and adapters ^17^. PCR amplicons were subjected to NGS (Novogene or HMS Biopolymers Facility). Computational barcode removal from raw NGS data was performed using in-house scripts. Low-read sgRNAs (i.e., sgRNAs represented by fewer than 10 reads in the plasmid library) were removed from the read count files. All subsequent read count and data analysis steps were performed using MAGeCK 0.5.9 ^77^.

### Gene set enrichment analysis (GSEA) of screen datasets

GSEA was performed using PANGEA ^27^. Prior to analyses, we updated PANGEA ^27^ to a PANGEA version 2 that adds coverage for *I. scapularis;* PANGEA v2 was used for all analyses. For *Drosophila* ortholog-based GSEAs, we used the full set of *Drosophila* orthologs of the *I. scapularis* genes screened for background correction. *I. scapularis* genes were mapped to *Drosophila* orthologs using the DIOPT mapping approach ^35^. The gene sets queried and other relevant information is annotated as metadata on the PANGEA output supplemental files.

### Analysis of human fitness gene datasets for cross-species comparison

Human essential gene data were obtained from the Dependency Map (DepMap) Public 26Q1 dataset (depmap.org; ^38^). The CRISPR-Cas9 gene essentiality scores (CERES algorithm) were downloaded using the depmap R package, encompassing 17,386 genes across 1,086 cell lines. Fitness genes were defined as those that had a CERES score of −0.5 in ≥95% of cell lines.

### Comparative analysis of Ctr1-family proteins

To compare Cup2 with functional domain and family annotations, a representative Cup2 amino acid sequence (NCBI XP_029847050.1) was inputted at the EBI-EMBL instance of InterProScan and analyzed using default settings ^44^. For alignment and phylogenetic tree construction, protein sequences from NCBI (*I. scapularis* Cup, XP_029847050.1; *I. scapularis* Ctr1A, XP_042148374.1; *D. melanogaster* Ctr1A, AFH07266.1; *D. melanogaster* Ctr1B, NP_649790.1; *D. melanogaster* Ctr1C, AAF57103.1; *Saccharomyces cerevisiae* CTR1, NP_015449.1; *S. cerevisiae* CTR2, NP_012045.3; *S. cerevisiae* CTR3, NP_013515.3; *Homo sapiens* SLC31A1, EAW87358.1, *H. sapiens* SLC31A2, EAW87355.1) were used as the input with default settings at the EMBL-EBI Job Dispatcher instance of Clustal Omega ^49^. The phylogenetic tree was enlarged proportionally using the graphical user interface at the EMBL-EBI Clustal Omega site prior to import into Adobe Illustrator.

### Cell competition assay in the presence of CuCl_2_

A total of 9 × 10⁶ RMCE+ Cas9 ISE18 DsRed cells were seeded into a T25 flask in 5 mL of tick medium. Cells were transfected with a plasmid mixture containing equimolar amounts of CAG-PhiC31 and an sgRNA donor library plasmid; either pLib025-iRFP720 (encoding sgRNAs targeting *Ctr1a or Cup2*, or a non-targeting sgRNA). Following transfection, cells were maintained for 25 days to allow stable integration and expression. Cells were not enriched for iRFP720, leaving a mixed population of RMCE+ Cas9+ ISE18 DsRed cells with and without the pLib025-iRFP720 integrated plasmid. After the 25-day integration period, a total of 9 × 10⁶ mixed cells were plated into T25 flasks under two conditions: untreated or treated with 60 μM CuCl₂. The initial mixed population was analyzed by flow cytometry. Cells were then cultured for 35 days with or without CuCl_2_, with medium changes and passaging performed every 4–6 days to maintain logarithmic growth. On day 35, cells were harvested and analyzed by flow cytometry to quantify the relative abundance of iRFP720⁺ populations.

### DA *in vivo* mortality assay in *I. scapularis* nymphs

*I. scapularis* nymphs were purchased from Oklahoma State University and the University of Minnesota. A stock solution of 10 mM of DA (Cayman Chemical) in DMSO was diluted in water to final concentrations of 5 or 1 µM. Aliquots of 100 µL of each dilution were then distributed in 1.5 mL microcentrifuge tubes. Control solution for each concentration contained equivalent amounts of DMSO. Twenty-five nymphs were transferred into each tube and centrifuged at maximum speed for 1 min to ensure contact with the solution. After 15 minutes of immersion, DA or control solutions were removed. A mesh-covered opening was created in the lid of each microcentrifuge tube to allow air exchange and maintain humidity. Ticks were then incubated in a Percival I-30BLL incubator at 23°C with 85% relative humidity and a 12/12 h light/dark photoperiod regimen. Ticks were treated again with DA or control solutions every other day and mortality was recorded over a period of 7 days.

### MTT assay of viability following RNAi knockdown and DA treatment

Small interfering RNA (siRNA) and scramble control (scRNA) for *PIG-A* were synthesized by Millipore Sigma with UU overhangs. A total of 3 x 10^6^ ISE6 cells were pelleted by centrifugation at 150 x g for 10 minutes at room temperature. The pellet was washed with 10mL of DPBS (Gibco) and resuspended in 30 µL of prewarmed nucleofection SF Buffer (Lonza Biosciences). Subsequently, 1.6 µg of siRNA or scRNA were added to the suspension and cells were subjected to the EN150 pulse condition using a 4D-Nucleofector system (Lonza Biosciences). Cells were incubated for 10 minutes at room temperature before adding pre-warmed L15C300 complete media. Cells were then seeded at 1 x 10^5^ cells/well in a 96-well plate. On day 3 and day 6 post-silencing, cells were exposed to 100, 50, 5 and 0.5 µM of DA (Cayman chemical). On day 9 post-silencing, cell metabolic activity was measured using CyQuant MTT cell viability assay kit (Invitrogen). Culture media of each well was replaced with fresh culture medium before adding 10 µL of 12-mM MTT stock solution and incubated at 34°C for 5 hours. After incubation, 85 µL of media was removed from each well and the resulting formazan crystals were solubilized with 50 µL of DMSO followed by incubation at 34°C for 10 minutes. Absorbance was read at 540 nm using iMark Microplate Absorbance Reader (Bio-Rad). Blank wells which consisted of MTT and medium alone were included to establish the baseline absorbance. Each condition was tested in quadruplicate.

### Confirmation of RNAi knockdown by quantitative real time PCR (qPCR)

Cells that were treated with the RNAi reagent but not exposed to DA were used to test knockdown efficiency by qPCR. Total RNA was extracted using a PureLink RNA Mini kit (Invitrogen) from samples preserved in TRIzol (Invitrogen). Next, cDNA synthesis was performed with a Verso cDNA Synthesis kit (Thermo Scientific) using 70-130 ng of RNA. To test knockdown efficiency, *pig-a* expression was measured by qPCR using 1 µL of cDNA and 2X Universal SYBR Green Fast qPCR Mix (ABclonal) in CFX96 Touch Real-Time PCR Detection System (Bio-rad). The qPCR conditions used were: (1) 95°C for 3 minutes; (2) 40 cycles of 95°C for 10 seconds, 54°C for 30 seconds; and (3) melting curve analysis to confirm the specificity of the reaction. No-template controls were added to the assay to verify absence of contamination. Reactions on each sample were run in duplicate. Gene expression was calculated by relative quantification normalized to *I. scapularis actin*.

## Data Availability

Transcriptomics datasets for ISE18, ISE6 and IDE2 are available at NCBI GEO (BioProject ID PRJNA1123289). Cell genome sequence datasets for ISE18 and ISE6 are available at NCBI SRA (BioProject ID SAMN57160449 for ISE18 and SAMN57160448 for ISE6). Genome-wide CRISPR-Cas9 sgRNA designs for *I. scapularis* can be searched, viewed, and downloaded at CRISPR GuideXpress (https://www.flyrnai.org/tools/fly2mosquito/web/) ^19^ and the sequences of sgRNAs included in this study are in the analyzed screen data files. PANGEA can be used to query public gene sets for *D. melanogaster* or human gene sets as reported previously ^27^, or to query public gene sets for *I. scapularis* as newly added for this work (the *I. scapularis* tab can be directly accessed at https://www.flyrnai.org/tools/pangea/web/home/6945). Analyzed screen datasets are included as supplemental files and raw data files are available upon request.

## Acknowledgements

We thank Ulrike Munderloh, Timothy Kurtti, and Nicole Burkhardt at the University of Minnesota for generously sharing cell lines and providing expert advice. We thank Isobel Ronai for helpful discussions. We thank the Harvard Medical School (HMS) MicRoN Imaging Core, the HMS Immunology Flow Cytometry Core, and the HMS Biopolymers Facility for expert advice and services. We also thank Elisabeth Brown and Richard Binari for exceptional lab support. Most of this work was completed at the Arthropod Cell Screening Facility (ACSF; formerly known as the Drosophila RNAi Screening Facility) in the Perrimon lab at HMS. We thank the staff of the ACSF/DRSC, past and present, as well as past and present members of the Perrimon lab, for advice and encouragement. BioRender (https://BioRender.com) was used to generate illustrations in Fig. 1 and Suppl. Fig. 2. This project received early seed funding from the Fairbairn Foundation HMS Lyme Initiative (NP, Principal Investigator; SEM and JHFP, Co-Investigators). This work was supported by the National Institute on Allergy and Infectious Disease (NIAID) at the U.S. National Institutes of Health (R21AI168592 to NP, Principal Investigator; SEM and JHFP, Co-Investigators). Additional support from NIAID R01AI186279 (NP, Principal Investigator; SEM and JHFP, Co-Investigators), NIAID R01AI116523 (JHFP, Principal Investigator) and NIAID P01AI138949 (JHFP, Multiple Principal Investigator). NP is an investigator of Howard Hughes Medical Institute.

## Open Access Statement

The article submitted together with this notice is subject to the Immediate Access to Research policy of the Howard Hughes Medical Institute (“HHMI”). In accordance with this policy: (i) a preprint of this article either has been, or will be, deposited on a preprint server under a Creative Commons Attribution 4.0 International (CC BY 4.0) license and (ii) an additional author-published revised version of this article incorporating peer review feedback and/or new results or analysis either has been, or prior to journal publication will be, deposited on a preprint server under a CC BY 4.0 license. In addition, a non-exclusive CC BY 4.0 license to this article has been granted to the public and HHMI has a sublicensable, non-exclusive license to this article.

THIS ARTICLE IS SUBMITTED FOR REVIEW AND ACCEPTANCE SUBJECT TO THESE PRE-EXISTING CONDITIONS, REQUIREMENTS AND LICENSES.

If you have any concerns with any of these pre-existing conditions, requirements or licenses, please contact the corresponding author immediately.

## Author Contributions

Conceptualization: MB, EM, JHFP, YH, RV, NP, SEM. Experimental investigation: MB, WM, SG, AWC, EM, AW, LQ, HV, EAL, HJLY, CA, NS. Reagent library design and addition of *I. scapularis* to PANGEA: AV and YH. Data analysis and visualization: MB, AWC, EM, AW, WC, YH, RV, SEM. Protein structure prediction: AK. Data management and deposition: MB, RV, YH, SEM. Project oversight and fundraising: MB, JHFP, SEM, NP. Initial writing and figure preparation: MB, WM, AWC, EM, AW, AK, YH, RV, SEM. Additional writing and revision: MB, WM, SG, AWC, EM, AW, LQ, AK, HV, EAL, HJLY, NS, JHFP, YH, RV, NP, SEM. All authors read and approved the manuscript.

## Materials List

An Excel file with tabs for cells, plasmids, and oligonucleotide primers is included as **Suppl. File 12**.

## Supplemental Figure Legends

**Supplemental Figure 1:**
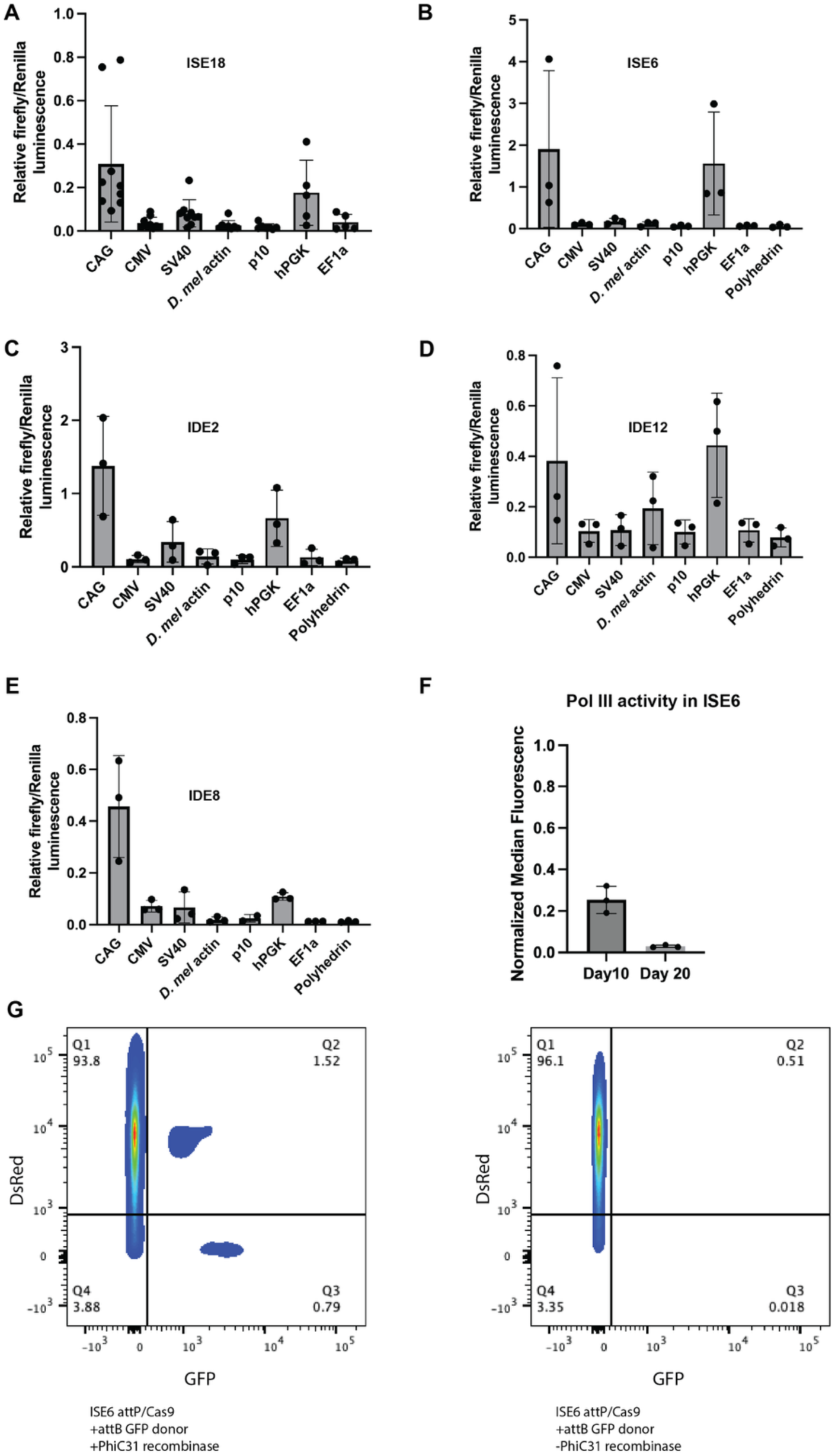
Molecular tool analyses in multiple *I. scapularis* cell lines. (**A**) – (E) Comparison of pol II promoters in five *I. scapularis* cell lines. Graphs of expression of firefly luciferase from the indicated pol II promoter normalized to CAG-Renilla luciferase. *D. mel* actin, *Drosophila melanogaster* Actin5c promoter. (F) Assay of pol III activity in RMCE+ Cas9+ ISE6 DsRed cells. Graph of DsRed levels following expression of a DsRed-targeting sgRNA with the 025 *I. scapularis* pol III promoter (only 025 was tested). (G) Evidence of recombination mediated cassette exchange (RMCE) in RMCE+ Cas9+ ISE6 DsRed cells. An attB-containing CAG-GFP cassette (pLib025-GFP) was transfected into RMCE+ Cas9+ ISE6 DsRed cells in the presence (left) or absence (right) of a plasmid expressing PhiC31 integrase. Shown are representative flow cytometry plots. GFP-positive cells are observed in the population with PhiC31 (left), indicating successful cassette exchange.

**Supplemental Figure 2:**
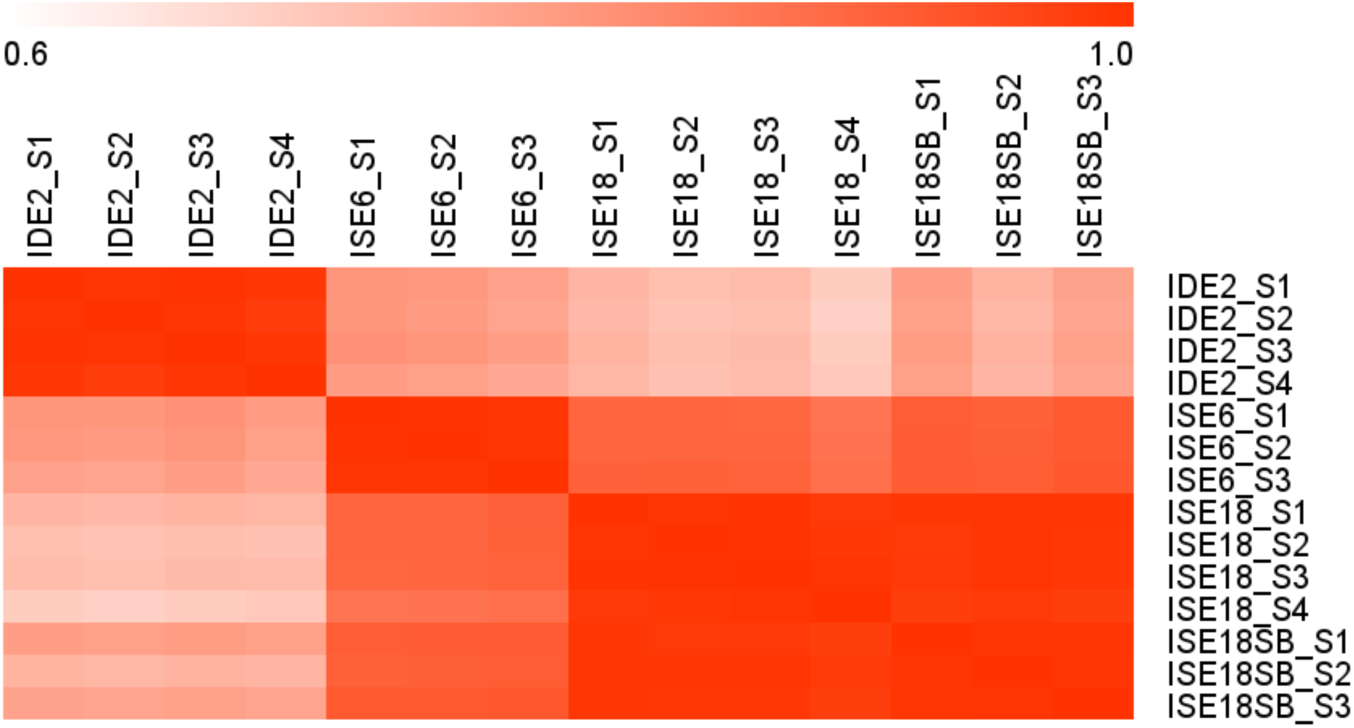
Comparison of expression patterns in *I. scapularis* cultured cell lines. Heatmap of the Pearson correlation for gene expression in IDE2, ISE6, ISE18, and a pool of RMCE+ ISE18 DsRed cells enriched for. Four (IDE2, ISE18) or three (ISE6, RMCE+ ISE18 DsRed) replicate samples are shown (S1, S2, etc.). Highest correlations are seen within replicate samples from the same cell line, and between ISE18 and RMCE+ ISE18 DsRed, and ISE6 and ISE18 are more similar to one another than either is to IDE2.

**Supplemental Figure 3:**
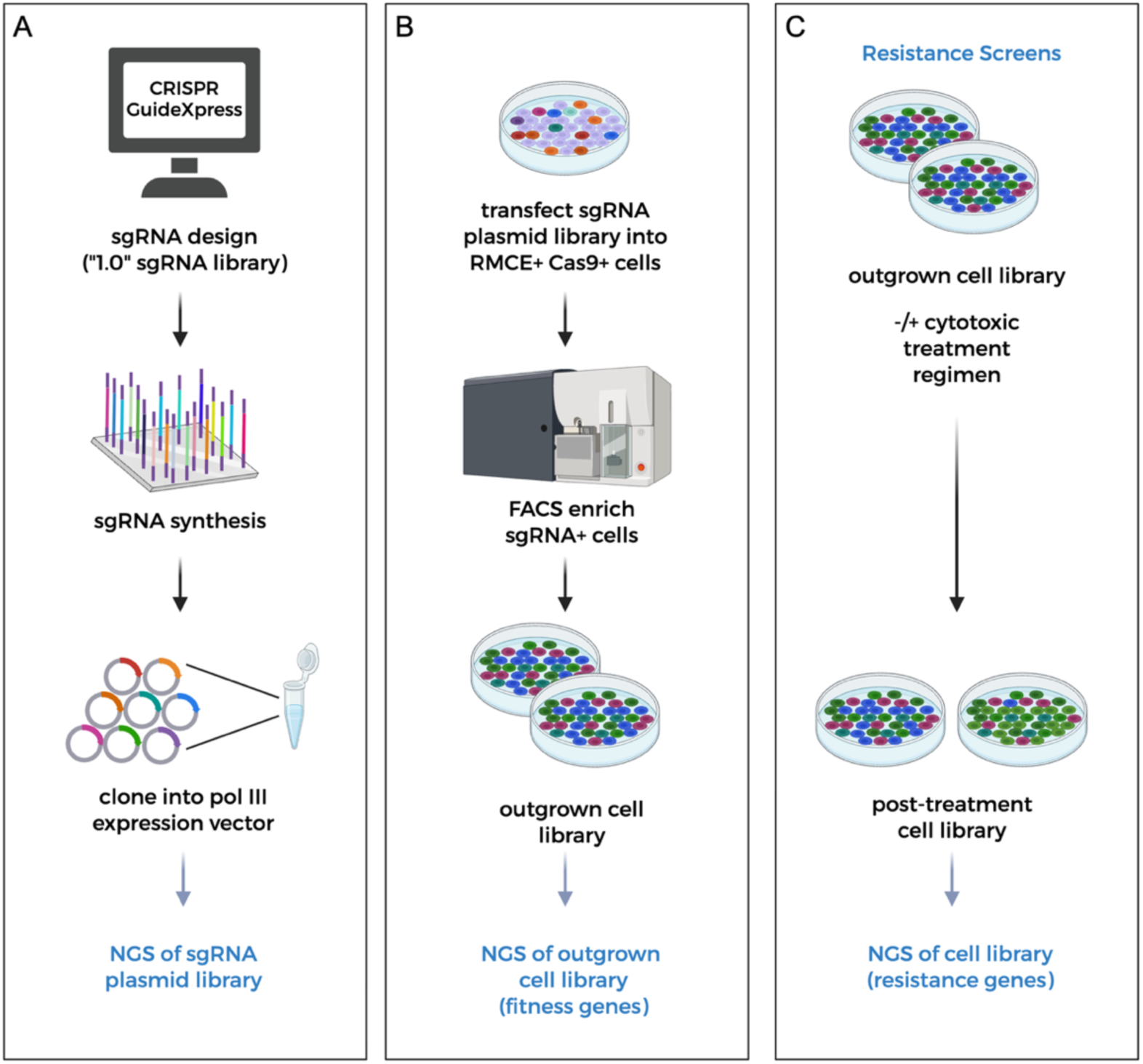
Summary of *I. scapularis* ISE18 CRISPR-Cas9 knockout (KO) cell screen workflows. (A) Single guide RNAs (sgRNAs) were designed using CRISPR GuideXpress, synthesized, and cloned into an *I. scapularis* pol III-driven sgRNA expression vector. (B) Cells were transfected with the sgRNA library and enriched for sgRNA integration via FACS. Samples of outgrown KO cell libraries were collected for the ‘drop-out’ (fitness) gene screen. (C) For treatment resistance screens (selection assays), cells from the FACS-enriched outgrown KO cell library were cultured in the absence (control) or presence of a cytotoxic treatment. The specific treatment regimen (e.g., timing of treatment, continuous treatment or cells were allowed to recover in standard media between treatments, etc.) differed for different screens. Following the treatment regimen, cells were collected for NGS.

**Supplemental Figure 4:**
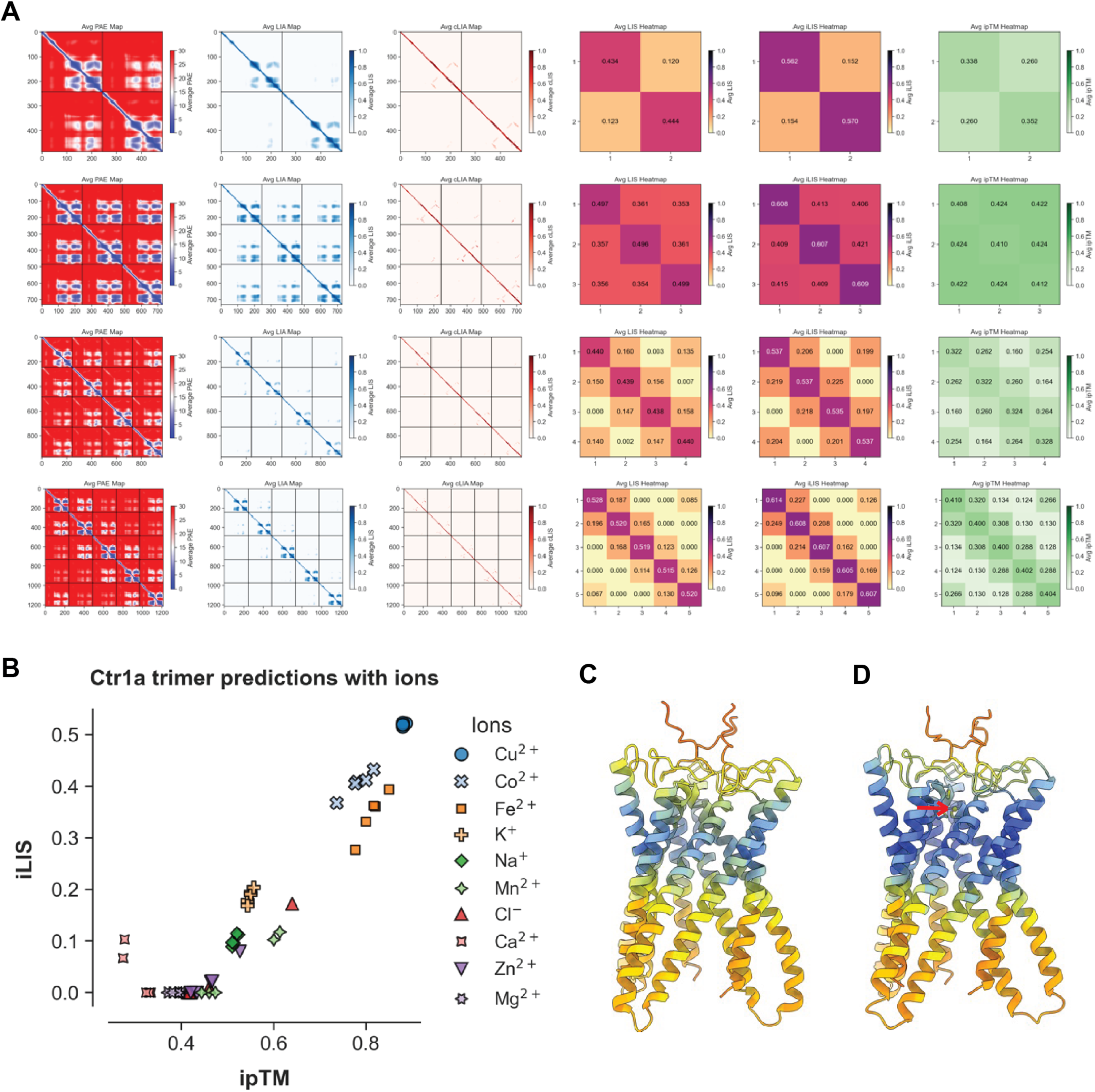
AlphaFold modeling of *I. scapularis* Ctr1A. (A) Evaluation of *I. scapularis* Ctr1A oligomeric states predicted by AlphaFold3. The panels display metrics averaged across five independent structural models for each tested subunit stoichiometry. Visualizations include the Predicted Aligned Error (PAE) map, Local Interaction Area (LIA) and contact-filtered LIA (cLIA) maps, and heatmaps for the Local Interaction Score (LIS), integrated LIS (iLIS), and interface predicted Template Modeling (ipTM) score. Low PAE (blue color) is suggestive of confident interactions. The trimeric assembly exhibits the highest iLIS and ipTM values, suggesting that it is the most structurally stable and confident oligomerization state. Plots generated as described in **Methods**. (B) AlphaFold3 predictions of Ctr1A interactions with different ions. The scatter plot compares the average iLIS against the ipTM score. Higher iLIS and ipTM values indicate a stronger predicted interaction and higher confidence in the protein-ion interface geometry, respectively. For each ion condition, five independent models were generated and plotted. The prediction with Cu^2+^ exhibits the strongest interaction among the tested ions. (C-D) AlphaFold3-predicted structure of the Ctr1a trimer alone (C) or in complex with Cu^2+^ (D). The structure is colored by pLDDT, which ranges from 0 to 100, where higher values (blue) indicate greater confidence in the local structure and lower values (orange) indicate lower confidence. The addition of Cu^2+^ increases the predicted rigidity of the trimer, as indicated by higher overall pLDDT scores. The red arrow points to the copper ion. The N-terminal region (residues 1–90) had a high PAE and was omitted for clarity.

**Supplemental Figure 5:**
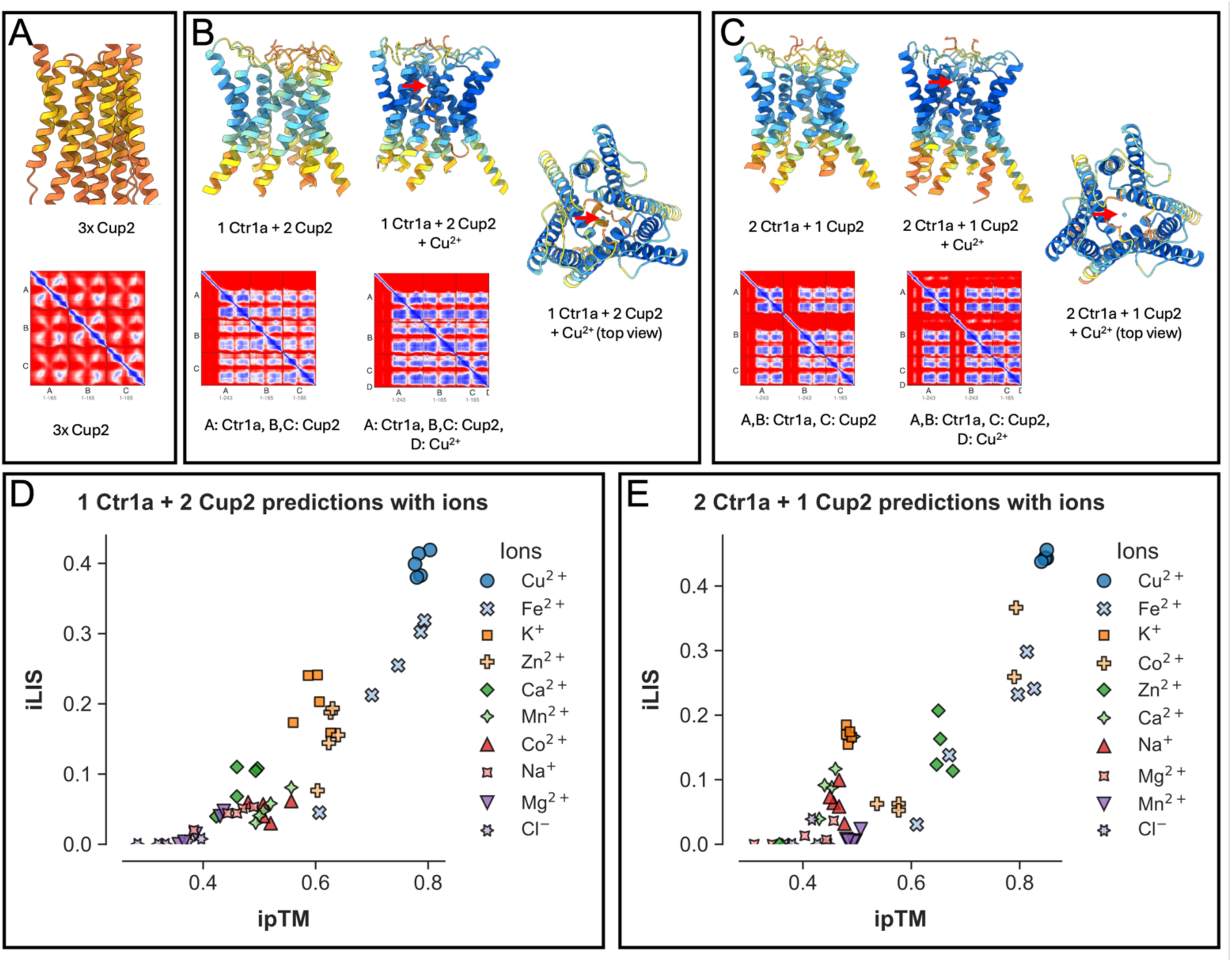
AlphaFold modeling of *I. scapularis* Cup2 and Cup2-Ctr1A heterotrimeric complexes. (A) Top, Cup2 homotrimer structure colored by pLDDT. Orange and yellow pLDDT indicate low-confidence predictions. Bottom, PAE map of the Cup2 homotrimer prediction. The low pLDDT and high PAE observed for the Cup2 homotrimer prediction suggests it is unlikely this structure forms. (B) Top, 1 x Ctr1A and 2 x Cup2 heterotrimer structure prediction colored by pLDDT. Compared to the Cup2 homotrimer, this heterotrimer has a higher pLDDT (sky blue), suggesting possible heterotrimer complex formation. Center, the same structure with a copper ion (red arrow). Note the darker blue (high-confidence in pLDDT, stable structure). Right, the same structure with a copper ion (red arrow), top view. Bottom, corresponding PAE maps for the heterotrimer without (left) or with (right) a copper ion. The heterotrimer with a copper ion shows lower PAE (darker blue), suggesting more stable complex (C) Top, 2 x Ctr1A and 1 x Cup2 heterotrimer structure prediction colored by pLDDT. Compared to the Cup2 homotrimer, this heterotrimer has a higher pLDDT (sky blue), suggesting a relatively stable structure. Center, the same structure with a copper ion (red arrow). Note the darker blue (high-confidence in pLDDT, stable structure). Right, the same structure with a copper ion (red arrow), top view. Bottom, corresponding PAE maps for the heterotrimer without (left) or with (right) a copper ion. The heterotrimer with a copper ion shows lower PAE (darker blue), suggesting a more stable complex. (E, F), Predicted ion preference for Ctr1A-Cup2 heterotrimers.

**Supplemental Figure 6:**
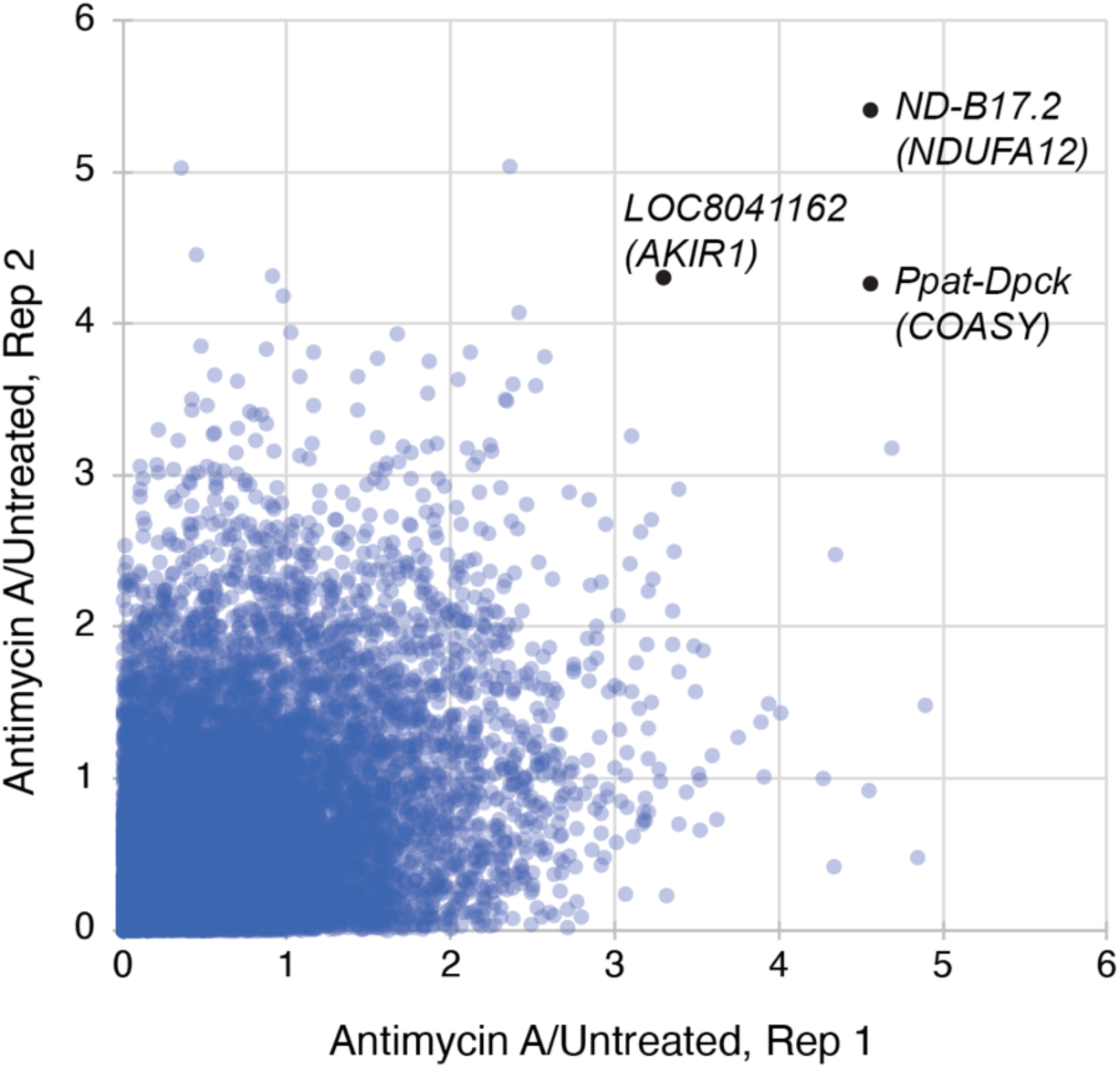
Genome-wide screen for resistance to Antimycin. **A.** Graph comparing data from the two replicate screens. *I. scapularis ND-B17.2* (NCBI Gene ID LOC8025722) is a predicted ortholog of human *NDUFA12*; *Ppat-Dpck* (LOC8025601), predicted ortholog of *COASY*; LOC8041162, predicted ortholog of *AKIR1*.

**Supplemental Figure 7:**
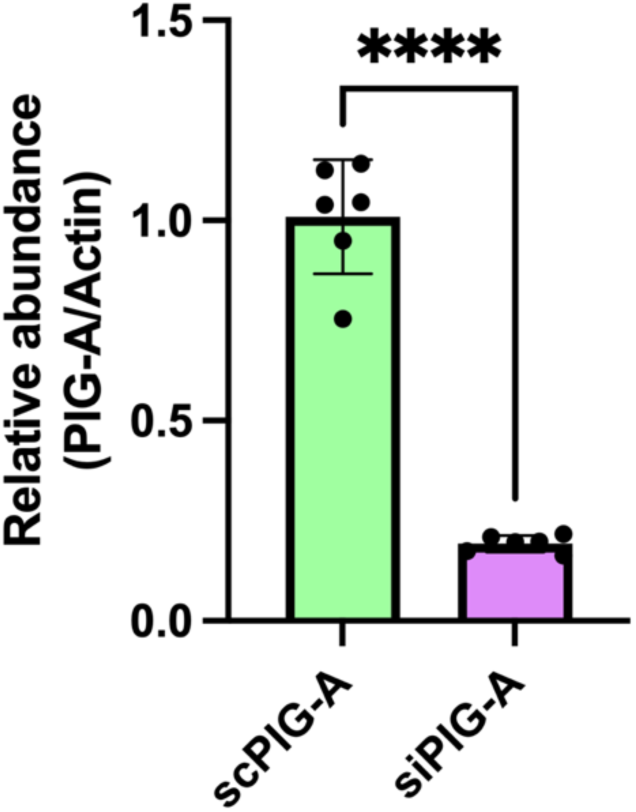
Quantitative real-time PCR (RT-qPCR) analysis of RNAi knockdown of *PIG-A* in ISE6 cells. Relative *PIG-A* mRNA expression was measured in ISE6 cells transfected with siRNAs targeting a non-targeting control siRNA (scPIG-A) or targeting the *PIG-A* gene (siPIG-A). Transcript levels were quantified by RT-qPCR and normalized using Actin as a reference gene.

## Supplemental Files

**Supplemental File 1**: Genome-wide *I. scapularis* CRISPR KO fitness screen dataset at sgRNA and gene levels annotated with gene expression and ortholog mapping.

**Supplemental File 2:** PANGEA GSEA with GO biological process and GO molecular function gene sets for the 850 *I. scapularis* fitness genes (FDR cutoff 1.0).

**Supplemental File 3:** PANGEA GSEAs with *Drosophila* gene sets for the *Drosophila* orthologs of the set of *I. scapularis* fitness genes (FDR cutoff 1.0).

**Supplemental File 4:** PANGEA GSEAs with *Drosophila* orthologs of the subset of *I. scapularis* fitness genes (FDR cutoff 0.1) that have *Drosophila* orthologs but those orthologs did not score as fitness genes in *Drosophila* S2R+ cells.

**Supplemental File 5:** Components of the TGF-beta/BMP signaling pathway are fitness genes in *I. scapularis* ISE18.

**Supplemental File 6:** PANGEA GSEAs with *I. scapularis* genes or their fly orthologs for growth-restrictive genes.

**Supplemental File 7:** Genome-wide *I. scapularis* CRISPR KO CuCl_2_ resistance screen dataset.

**Supplemental File 8:** PANGEA GSEAs with *I. scapularis* genes for which KO confers resistance to treatment with CuCl_2_.

**Supplemental File 9:** Genome-wide *I. scapularis* CRISPR KO Antimycin A resistance screen dataset.

**Supplemental File 10:** Genome-wide *I. scapularis* CRISPR KO Destruxin A (DA) resistance screen dataset.

**Supplemental File 11:** PANGEA GSEAs with *I. scapularis* genes for which KO confers resistance to treatment with Destruxin A (DA).

**Supplemental File 12:** Materials (cell lines, plasmids, oligonucleotide primers).

